# The influence of *GC*-biased gene conversion on nonadaptive sequence evolution in short introns of *Drosophila melanogaster*

**DOI:** 10.1101/2023.06.14.544886

**Authors:** Burçin Yıldırım, Claus Vogl

## Abstract

Population genetic inference of selection on the nucleotide sequence level often proceeds by comparison to a reference sequence evolving only under mutation and population demography. Among the few candidates for such a reference sequence is the 5’ part of short introns (5SI) in *Drosophila*. In addition to mutation and population demography, however, there is evidence for a weak force favoring *GC* bases, likely due to *GC*-biased gene conversion (gBGC), and for the effect of linked selection. Here, we use polymorphism and divergence data of *Drosophila melanogaster* to detect and describe the forces affecting the evolution of the 5SI. We separately analyze mutation classes, compare them between chromosomes, and relate them to recombination rate frequencies. *GC*-conservative mutations seem to be mainly influenced by mutation and drift, with linked selection mostly causing differences between the central and the peripheral (*i*.*e*., telomeric and centromeric) regions of the chromosome arms. Comparing *GC*-conservative mutation patterns between autosomes and the X chromosome, showed differences in mutation rates, rather than linked selection, in the central chromosomal regions after accounting for differences in effective populations sizes. On the other hand, *GC*-changing mutations show asymmetric site frequency spectra, indicating the presence of gBGC, varying among mutation classes and in intensity along chromosomes, but approximately equal in strength in autosomes and the X chromosome.

## Introduction

DNA sequence polymorphism and divergence data have been used to infer evolutionary processes in many fields of evolutionary biology, from molecular evolution to anthropology. Whether the aim is phylogenetic reconstruction, demographic inference, or detection of selection, it is difficult to tease apart the different population genetic forces that determine DNA sequence patterns. According to the neutral theory of molecular evolution (Kimura, 1983), the majority of mutations are selectively neutral, subject only to mutation and random drift, or strongly deleterious, while positively selected mutations play a minor role. In this theory, purifying selection efficiently removes deleterious mutations, and positively selected mutations rapidly reach fixation, hence they do not contribute to segregating variation. Early analysis methods often tested for deviation from neutral equilibrium (*e*.*g*., Tajima, 1989; Fu & Li, 1993). However, with the availability of genome-wide data and advanced analytical methods, it has become evident that sequences are rarely at equilibrium (Thornton et al., 2007). Furthermore, the indirect effect of selection on linked neutral sites, via background selection or selective sweeps, has been documented (Smith & Haigh, 1974; Charlesworth et al., 1993; Schrider et al., 2016). In light of these observations, modern analysis methods strive to adapt neutral models to incorporate evolutionary processes that are common to the genome (Johri et al., 2020, 2022).

Sequence classes that are known to evolve under nonadaptive forces are valuable resources for constructing such neutral models. Understanding the evolutionary processes that act on these sequences is crucial for comprehending the dynamics of genome evolution. Furthermore, they can be used as a reference in population genetics inference. Failing to account for a specific evolutionary force or its interaction with other forces can lead to biased inferences (Lartillot, 2012; Bolívar et al., 2016, 2018; Galtier et al., 2018; Borges et al., 2019; Bolívar et al., 2019; Boman et al., 2021). Hence, it is important to consider the relative contributions of various population genetic forces, which may vary depending on the organism under study and among different genomic regions.

Fourfold degenerate sites, or generally synonymous sites, have been regarded as neutrally evolving, because an exchange of a nucleotide does not affect the encoded amino acid. Although all genomes show some degree of codon usage bias in these sites (Hershberg & Petrov, 2008), the intensity of selection has been generally considered weak in relation to drift. Therefore, such sites have been used as a neutral reference for many tests (*e*.*g*., McDonald & Kreitman, 1991; Yang et al., 2000). These tests assume reduction in polymorphism and divergence due to purging of deleterious mutant alleles with directional selection. This assumption is a consequence of the infinite sites model, where mutations are considered irreversible (Kimura, 1969). With reversible mutations, however, weak selection opposing the mutation bias may increase polymorphism and divergence levels compared to neutrality (McVean & Charlesworth, 1999; Vogl & Mikula, 2021), creating unexpected patterns. An additional challenge for using fourfold degenerate sites as a neutral reference comes from studies showing that codon usage bias may vary from very weak to strong depending on the species, the codon and the position in the genome (Chamary et al., 2006; Lawrie et al., 2013). Thus, it is not clear *a priori* which codon is favored for a specific site in a specific species, and preferences may evolve.

A similarly weak force is *GC*-biased gene conversion (gBGC), which is tied to the repair of double-strand breaks (DSBs) by homologous recombination. When a site is heterozygous for a strong (*S*; *G* and *C* bases) and a weak (*W*; *A* and *T* bases) allele, the heteroduplex mismatch formed during the DSB repair might resolve biased towards the strong base (Marais, 2003). The biased resolution of these *GC*-changing mutations (*S*↔ *W*) leads to a nonadaptive directional force, indistinguishable in its effect from selection for *GC* bases (Duret & Galtier, 2009; Bolívar et al., 2018). At the molecular level, the repair efficiency of *GC*-changing mutations might change between transitions and transversions (Dohet et al., 1985; Holmes et al., 1990), and the effects of it might also be observed in the gBGC dynamics at the population genetic level (Lartillot, 2012; Bergman & Schierup, 2021). Studies have shown that failing to account for gBGC may lead to biased inference of selection and demography (Bolívar et al., 2018; Pouyet et al., 2018; Galtier et al., 2018). Additionally, the effects of gBGC on diversity may be falsely interpreted as a consequence of linked selection or Hill-Robertson interference (Bolívar et al., 2016; Boman et al., 2021) and confound the detection of expectations from nearly neutral theory (Bolívar et al., 2019) due to its relationship with recombination.

In flies of the genus *Drosophila*, it is known that a large fraction of the genome, including intronic and intergenic noncoding sequences, is under selective constraints (Andolfatto, 2005; Haddrill & Charlesworth, 2008; Haddrill et al., 2008). Synonymous codons are also subject to selection due to codon usage bias (Akashi, 1994, 1995) with variable intensity depending on species and codons (Singh et al., 2009; Zeng, 2010). So far, the best candidates for neutrally evolving, unconstrained sites in *Drosophila* are the nucleotides at position 8-30 on the 5’ end of introns shorter than 65 bp (hereafter 5SI) (Halligan & Keightley, 2006; Parsch et al., 2010; Clemente & Vogl, 2012; Yıldırım & Vogl, 2023). These sites exhibit higher divergence and polymorphism levels compared to other regions in introns (Parsch et al., 2010). Longer introns contain more functional elements, as shown by a negative correlation between divergence and length (Haddrill et al., 2005) and most other sequences inside short introns are likely under selection due to their association with splicing (Yıldırım & Vogl, 2023). Thus, many studieswere based on the premise that 5SI sequences evolve neutrally and therefore can be used to infer directional selection on fourfold degenerate sites (Lawrie et al., 2013; Machado et al., 2020) or demography before detecting sweep signatures (Garud et al., 2015). Indeed, the premise of this study is also that sequences in the 5SI evolve neutrally.

Despite the utility of the 5SI as a neutral reference in population genetic analyses, biallelic frequency spectra of weak vs strong bases deviate from the neutral prediction of symmetry, with an excess of high-frequency *GC* variants (Clemente & Vogl, 2012; Jackson et al., 2017; Jackson & Charlesworth, 2021), which indicates the presence of a directional force, likely gBGC. The presence of gBGC in the fruit fly *Drosophila* has been long debated and remained inconclusive. Clemente & Vogl (2012) explained the asymmetry in site-frequency spectra (SFS) of *AT* -to-*GC* polymorphism in *D. melanogaster* by a shift in mutation bias towards *AT* and a context-dependent mutational pattern. Robinson et al. (2014) claimed that the effect of gBGC is unlikely to impact genome evolution patterns. Other studies showed evidence for gBGC, but differed in their claims on which chromosome and in which species it operates (Haddrill & Charlesworth, 2008; de Procé et al., 2012; Jackson et al., 2017).

Most recently, Jackson & Charlesworth (2021) demonstrated the existence of a *GC*-favoring directional force in autosomal 5SI sites of both *D. simulans* and *D. melanogaster*, using unfolded SFS of *GC*-changing mutations (*S*→ *W, W*→ *S*) from a larger population data set and better reference genomes than the previous studies. This immediately raises interesting research questions: Does gBGC affect transitions and transversions similarly? Is gBGC also present in the X chromosome of *Drosophila*? If the pattern of gBGC is recombination-dependent, we expect about equal effects on autosomes and the X chromosome in *Drosophila*, as there is no recombination in males. Do we therefore find similar effects of gBGC on the X chromosome? Most importantly, it is known from other species that the presence of gBGC can lead to biased interpretations of the effects of linked selection and other nonadaptive forces. Variation in polymorphism along chromosomes that has been shown to be correlated with recombination rate variation in *Drosophila* is accepted to be the result of linked selection (Begun & Aquadro, 1992), but may also be affected by gBGC. How does the previously neglected presence of gBGC in *Drosophila* influence our interpretation of neutral sequence evolution patterns in autosomes and the X chromosome? Are patterns that have been attributed to linked selection affected or actually caused by gBGC?

In this study, we address these questions by utilizing 5SI from one of the largest and most accurate polymorphism data from the ancestral population of *D. melanogaster* and divergence data with *D. simulans* (Rogers et al., 2014; Lack et al., 2015; Jackson et al., 2017). The divergence time between these two species is conveniently so small that very few double mutations separating the species are expected, but large enough that little shared polymorphism is expected (estimates are given in the Results section). Therefore we can assume that only single mutations give rise to polymorphic and divergent sites. This allows us to compare estimates of polymorphism and divergence among different mutation classes and between autosomes and the X chromosome to identify and describe the relative contributions of various nonadaptive population genetic forces, such as gBGC, mutation, and drift, in shaping the evolution of this neutral sequence class. Specifically, we contrast *GC*-conservative mutation classes, which are expected to be unaffected by gBGC, with *GC*-changing mutations to answer the following questions: What is the relative effect of linked selection and gBGC on the variation in polymorphism and divergence i) along chromosomes and ii) between autosomes and X chromosomes? iii) How does variation in recombination rates modulate these effects? While addressing these questions, we also provide a detailed description of the dynamics of gBGC separately for transitions and transversions, both along and between chromosomes.

## Materials and Methods

### Data Used in the Analyses

We analyzed previously published whole-genome data from a population of *D. melanogaster* from the ancestral range in Zambia (Lack et al., 2015). After excluding individuals showing admixture with European populations (Lack et al., 2015), the dataset consists of 69 individuals both for autosomes and X chromosome. Sequences were obtained as consensus FASTA files, and the full description of the data processing (sampling, sequencing, variant calling) can be found in Lack et al. (2015). We also obtained consensus FASTA files from a population sample of *Drosophila simulans* from Madagascar including 21 individuals for all chromosomes (Rogers et al., 2014; Jackson et al., 2017).

Using annotations from the reference genomes of *D. melanogaster* (r5.57 from http://www.flybase.org/) and *D. simulans* (Hu et al., 2013), orthologous intron coordinates were extracted and alignments of all samples were created. To avoid including the same intron sequence more than once due to alternatively spliced isoforms annotated the GFF file (see https://www.ensembl.org/info/website/upload/gff.html; last accessed November 1, 2020), only one entry of introns with overlapping coordinates were used. For this, we chose introns belonging to the longest transcript of a gene. Bases in position 8-30 in short introns (≤ 65 bp) were extracted to use as a proxy of least constrained sites (Halligan & Keightley, 2006; Parsch et al., 2010). The analyses described here and below were performed using custom R and shell scripts.

### Polymorphism and divergence estimates

We inferred site frequency spectra from the Zambian *D. melanogaster* samples for all six possible combinations of base pairs, for both autosomal and X-linked short introns. We filtered out sites that overlapped coding sequences, contained an undefined nucleotide state in at least one of the sequences in the sample alignment and sites with more than two alleles. We note that sites with more than two alleles make up only a proportion of 5 ·10^−3^ of the polymorphic sites, are largely attributed to technical errors, and therefore usually filtered out (*e*.*g*., Bergman et al., 2017; Jackson & Charlesworth, 2021). We thus expect this filtering to negligibly affect our analyses.

The scaled mutation rate, denoted as *θ*, is the product of mutation rate per site per generation (*μ*) and effective population size (*N*_*e*_). An estimator of this scaled mutation rate, the Ewens-Watterson estimator (*θ*_*W*_), is defined as *L*_*p*_*/LH*_*M*−1_, where *L* is the total number of sites, *L*_*p*_ is the number of polymorphic sites, *M* is the sample size, and *H* is the harmonic number, given by the formula 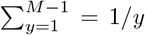 (Ewens, 1974; Watterson, 1975). Using effectively neutral sites, *θ* is estimated to be less than 10^−2^ in eukaryotes, and mutation rates decreases from 10^−8^ to 10^−10^ with increasing effective population size (Lynch et al., 2016). Therefore, methods generally consider small *θ* approximations (*e*.*g*., Vogl & Bergman, 2015; Burden & Tang, 2016).

We estimate scaled mutation rates under a multi-allelic model to infer the complete 4 *×* 4 mutation rate matrix from allele frequency data. We also assume mutation rates are low, such that segregation of more than two alleles in the population is negligible. Considering *i* and *j* stand for the four bases *i, j ϵ* (*A, T, G, C*), this implies 12 parameters, *θ*_*ij*_ = 4*N*_*e*_*μ*_*ij*_. We further assume strand symmetric mutation, where nucleotides *A* and *T* or *G* and *C* are interchangeable. This reduces the number of parameters from twelve to six and allows the use of the maximum likelihood estimators (MLEs) from Vogl et al. (2020). With SFS data available from *L* loci with *M* genomes, let *L*_*ij*_ be the counts of polymorphic sites and *L*_*i*_ the counts of monomorphic sites. The six MLEs for the scaled mutation rates are variants of Ewens-Watterson estimator (*θ*_*W*_) under multi-allelic model and defined as:

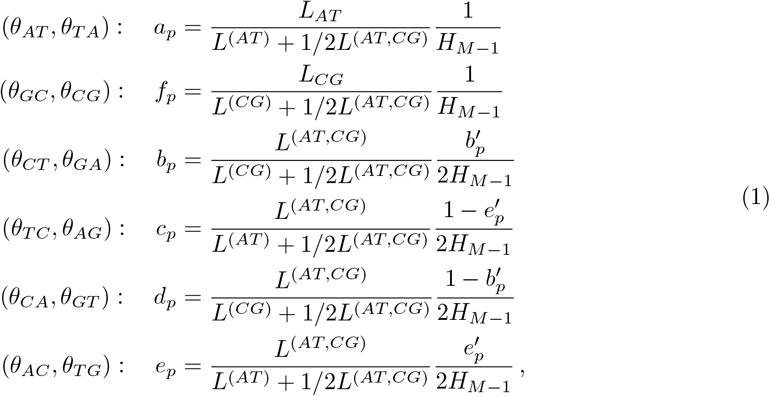

Where *L*^*(AT)*^ = *L*_*A*_ + *L*_*T*_ + *L*_*AT*,_ *L*^*(CG)*^ = *L*_*C*_ + *L*_*G*_ + *L*_*CG*_ and *L*^*(AT,CG)*^ = *L*_*TC*_ + *L*_*TG*_. The parameters 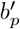 and 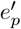 correspond to 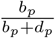 and 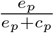, respectively. They are obtained by maximizing the following log likelihood:

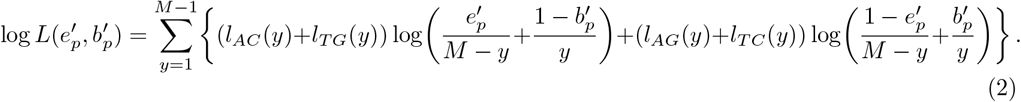

From polymorphism data, we also calculated mutation bias towards *AT* (*β*) as follows;

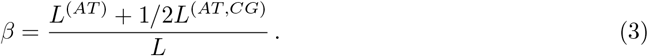

As a measure of divergence between *D. melanogaster* and *D. simulans*, we used an estimate corresponding to *D*_*a*_, the net nucleotide differences between populations since the species split (Nei & Li, 1979). It is defined as *D*_*a*_ = *D*_*xy*_ − (*π*_*x*_ + *π*_*y*_)*/*2, where the expectation of the pairwise difference *D*_*xy*_ is 2*μt* + *θ*_*anc*_. *D*_*a*_ uses the average current levels of polymorphisms ((*π*_*x*_ + *π*_*y*_)*/*2) as a measure of ancestral polymorphism (*θ*_*anc*_), and subtracts this value from the total divergence to get an estimate of *δ* = 2*μt*, where *t* is the time in generations since the species split. If linked selection was present before the split of the two populations, in the ancestral population, *D*_*a*_ should be affected only slightly, while *D*_*xy*_ should be reduced (figure 1). Begun et al. (2007) showed that current targets of selection in *Drosophila* are also targets of recurrent selection, thus using a *D*_*a*_-like measure for divergence would minimize the effect of linked selection. We calculated such a divergence estimate from the alignment of two *D. melanogaster* and two *D. simulans* samples. Corresponding to the estimators of scaled mutation rates, six estimators of divergence were obtained for pairs of *δ*_*ij*_ = 2*μ*_*ij*_*t*:

**Figure 1:**
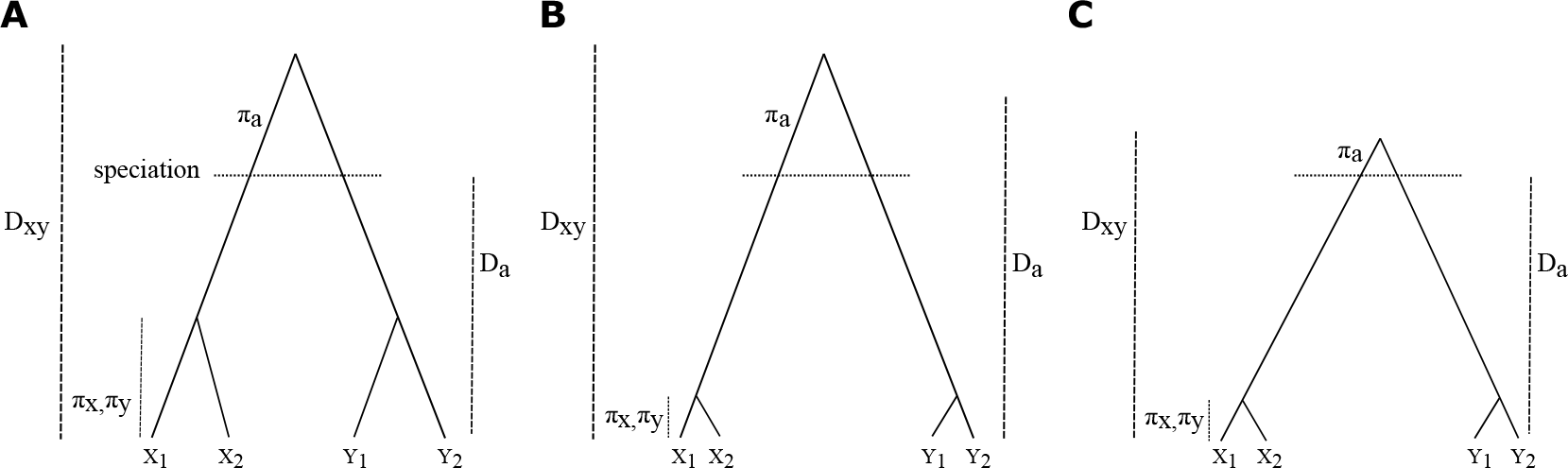
Genealogies of four samples (*X*_1_, *X*_2_, *Y*_1_, *Y*_2_) from two populations or species (X and Y) and measures of divergence with (B, C) and without (A) the effect of linked selection. *π*_*x*_, *π*_*y*_ and *π*_*a*_ represent the polymorphism level in current (X, Y) and ancestral populations, respectively. *D*_*xy*_ is a measure of divergence defined as average number of pairwise differences between sequences of populations. Lastly, *D*_*a*_ is a measure of pairwise differences since the split of populations. Panel A represents a scenario without linked selection, panel B represents the linked selection affecting current-day populations and panel C shows a scenario with recurrent linked selection affecting both ancestral and extant populations. Figures are adapted from Cruickshank & Hahn (2014).

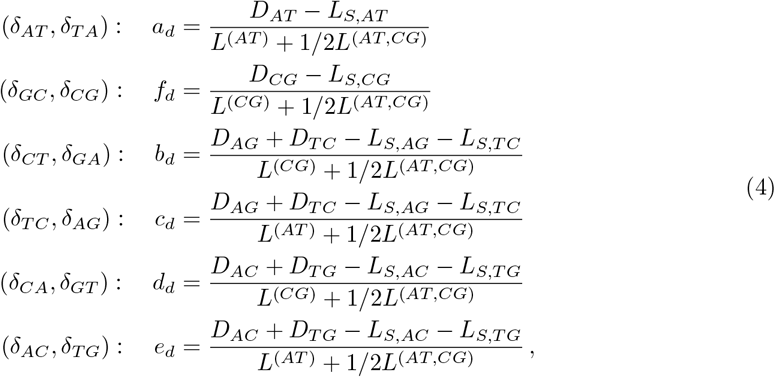

where *D*_*ij*_ corresponds to the number of sites differentially fixed for *i* and *j* nucleotides and *L*_*S,ij*_ to the number of ancestral shared polymorphism segregating for *i* and *j* in *D*.*melanogaster* and *D. simulans*.

Under equilibrium conditions, we note that for autosomes the expected neutral divergence E[*a*_*d*_] and the expected neutral scaled mutation rate or polymorphism E[*a*_*p*_] are proportionally affected by the same pair of mutation rates *μ*_*AT*_ and *μ*_*TA*_, such that we have:

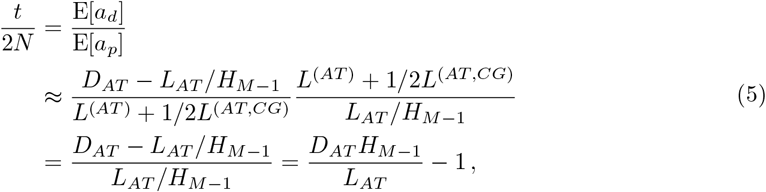

and similar for other pairs of polymorphism and divergence estimators. Assuming equality of male and female effective population sizes and accounting for the hemizygosity of males, the ratio of divergence over polymorphism for the autosomes would be 3*/*4 of the X chromosome:

(6) When a mutation class without a directional force is compared to another with a directional force *γ*, the “neutrality index” can be calculated as a ratio of ratios of divergence and polymorphism between the two classes assuming neutrality. Importantly, the mutational terms cancel when the ratio is taken. Let *r*_*N*_ stand for the ratio of expected divergence to expected polymorphism of the neutral class and *r*_*γ*_ for that with a directional force. Then *r*_*N*_ */r*_*γ*_ = ((*e*^*γ*^*/*2 − *e*^−*γ/*2^)*/γ*)^2^ (see equation 54 in Vogl & Mikula (2021)), which is always greater than 1 with *γ* ≠ 0 in equilibrium.

Due to filtering out the non-biallelic sites and missing polymorphism in *D. melanogaster* population data, different numbers of sites were available for polymorphism and divergence based analyses. To make analyses comparable, we included in the final dataset only the common sites, of which 137,699 were autosomal and 16,873 X chromosomal. We also analyzed polymorphism and divergence data from *D. simulans*, resulting in 150,613 autosomal and 19,412 X chromosomal common sites. The 95% confidence intervals for each point estimate were determined from 1000 bootstrap-resamples (Efron, 1979). Our estimates were calculated for whole chromosomes, as well as for the central and peripheral (telomeres and centromeres combined) regions of chromosome arms. Genomic locations for the central and peripheral regions were obtained from Comeron et al. (2012) (see their Table 3), which were defined according to the visibly reduced crossover rates in telomeres and centromeres. Unless otherwise stated, our reported estimates come from the central part of the chromosome arms after excluding telomeres and centromeres.

### Recombination rates and base composition

To study the co-variation of the inferred parameter estimates and recombination rates, we retrieved the recombination rate estimates, based on crossover events, from the *D. melanogaster* recombination map of Comeron et al. (2012). We divided the dataset into bins with approximately equal number of observations, before and after excluding peripheral regions of the chromosome arms with low recombination rates. Mean recombination rate for each bin is given in table S1. The site frequency spectra inferred separately for each recombination bin were used to estimate the polymorphism, divergence and the strength of gBGC.

We examined the base composition by looking at *GC* content, calculated for each intron separately from the reference genomes of *D. melanogaster* and *D. simulans*. The *GC* content of the four-fold degenerate sites (FFDS) from the same genes that introns located were also obtained and used as a proxy for background base composition. The *GC* content was determined as the number of *G* and *C* nucleotides divided by the total number of defined nucleotides, *i*.*e*., excluding undefined (*N*) nucleotides. To investigate the variation of *GC*-biased gene conversion with the base composition, we divided the dataset into five bins with approximately equal number of observations depending on the background *GC* content (*i*.*e*., FFDS *GC* content). The range of *GC* content for each bin is given in tables S2 and S3 for *D. melanogaster* and *D. simulans*, respectively.

### Estimates of gBGC strength

Heteroduplex mismatches formed during the repair of double strand breaks can involve either pairing between the bases *G* and *C* (strong: S:S), *A* and *T* (weak: W:W) or between strong and weak bases, S:W. Preferential resolution of the S:W mismatches into *G* : *C* rather than *A* : *T* leads to *GC*-biased gene conversion (gBGC) (Marais, 2003). Since gBGC only affects S:W mismatches they are referred to as *GC*-changing, while the others (S:S, W:W) are called *GC*-conservative. This categorization allows us to use unpolarized data to estimate *GC* bias while considering the site frequency spectra (SFS) of all six possible nucleotide pairs (Borges et al., 2019). gBGC is a directional force quantified by *B* = 4*N*_*e*_*b, b* is the conversion bias that depends on recombination rate, tract length and repair bias towards *GC* (Nagylaki, 1983).

We inferred the strength of this directional force, gBGC, under mutation-drift-directional force equilibrium by using the maximum likelihood estimator of Vogl & Bergman (2015). Assuming low mutation rate and binomial sampling, the probability of a mutation segregating at frequency *y* in the limit of large M is;

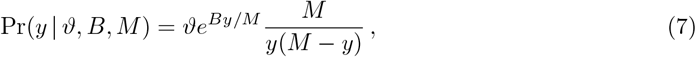

where *ϑ* = ((1 − *β*)*βθ*)*/*((1 − *β*)*e*^*B*^ + *β*) and *β* is mutation bias towards *AT*. The likelihood of the polymorphic loci is sufficient for the inference of *B* and expressed as:

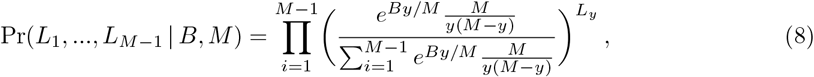

where *L*_*y*_ is the number of sites with *y GC*-changing mutations. *B* estimates were obtained by maximizing the likelihood in equation 8 for the SFS from *GC*-changing polymorphisms (*A/C, A/G, T/C* and *T/G*) of the 5SI. The SFS from *GC*-conservative polymorphisms (*A/T, G/C*) was considered as putatively neutral control (*B* = 0). We performed likelihood-ratio tests (LRT) to compare between the different nested models. Conditional on *B*, we also estimated the mutation bias towards 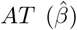. For the inference, the mutation bias parameter was set to *ϱ* = 1 − ((1 − *β*)*e*^*B*^)*/*((1 − *β*)*e*^*B*^ + *β*) with the maximum likelihood estimate of;

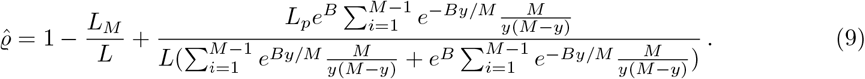

Given the estimate of 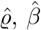 was recovered using 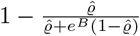.

We also inferred *B* while correcting for the effect of demography and population structure by introducing noise parameters *r*_*y*_ to the likelihood (equation 8)(Bergman & Schierup, 2021). We obtained *r*_*y*_ by comparing the neutral expectation of the SFS with the empirical SFS of the neutral sites, *i*.*e*., SFS of *GC*-conservative polymorphisms.

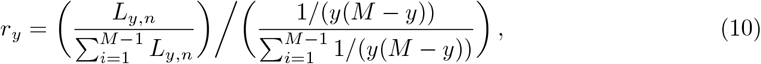

where *L*_*y,n*_ is the number of sites with *y GC*-conservative polymorphisms. Due to the large sample sizes *M* and splitting of the data according to mutation classes and genomic regions, counts of *r*_*y*_ = 0 are possible, which leads to undefined values when calculating the log-likelihood. To avoid that, we added pseudo-counts proportional to the equilibrium expectations 1*/y* + 1*/*(*M − y*). We note that accounting for demography in this way lowers the statistical power. The 95% confidence intervals for each estimate were constructed using a likelihood ratio test (Zhou, 2015).

## Results

We investigated the pattern of molecular evolution and variation in neutral short introns of *Drosophila melanogaster* and *D. simulans* by comparing different nucleotide classes along chromosomes, between chromosomes and in relation to recombination rates and *GC*-content to estimate the contribution of different nonadaptive forces to the observed patterns. We used both polymorphism and divergence data to get estimates of scaled mutation rates (or polymorphism or diversity, 4*N*_*e*_*μ*_*ij*_) and divergence (2*μ*_*ij*_*t*) for each nucleotide class. Instead of a bi-allelic mutation-drift model, we used a multi-allelic model, corresponding to the four bases, that can provide information about the possible differences in mutational bias and rate between different alleles. Given the four bases, this would imply 4 ·3 = 12 parameters.

If there is no DNA strand-specificity of mutation rates, the equal proportion of complementary bases (*i*.*e*., of the weak bases *A* and *T* and the strong bases *C* and *G*, respectively) along the DNA leads to strand symmetry, *i*.*e*., Chargaff’s second parity rule (Mitchell & Bridge, 2006). The unselected intronic sequences we analyzed exhibit only very minor deviations from strand symmetry (slight biases towards *T* and *C* over *A* and *G*, respectively) (Bergman et al., 2017), which are attributed to neutral processes, such as transcription-coupled asymmetries (Touchon et al., 2004). Thus, we obtained the MLE of scaled mutation rates by assuming neutral equilibrium and strand-symmetric mutation (Vogl et al., 2020), which reduced the number of parameters from twelve to six (*a*_*p*_, *b*_*p*_, *c*_*p*_, *d*_*p*_, *e*_*p*_, *f*_*p*_), see equation (1). Our inference further assumes that the scaled mutation rates are small, specifically, they should be below 0.05 or, more stringently, below 0.02 (Vogl & Clemente, 2012). This indeed holds true as the highest estimate of theta is approximately 0.024 (table 1). Therefore segregation of more than two alleles in the sample is negligible.

**Table 1:**
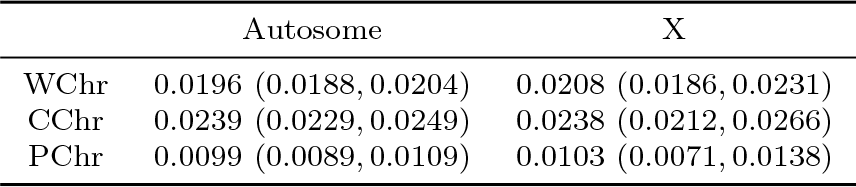
The overall expected heterozygosity estimates for different parts of the autosomes and X-chromosome (95% CIs in brackets). WChr: Whole chromosome, CChr: Central regions of the chromosome arms, PChr: Peripheral regions of the chromosome arms.

|  | Autosome | X |
| --- | --- | --- |
| WChr | 0.0196 (0.0188, 0.0204) | 0.0208 (0.0186, 0.0231) |
| CChr | 0.0239 (0.0229, 0.0249) | 0.0238 (0.0212, 0.0266) |
| PChr | 0.0099 (0.0089, 0.0109) | 0.0103 (0.0071, 0.0138) |

In addition to mutational differences, some of these nucleotide classes might be affected distinctly by other evolutionary forces. The *A* : *T* (*a*_*p*_) and *C* : *G* (*f*_*p*_) polymorphisms are transversions that exhibit negligible mutation bias in *Drosophila* and, more importantly, these two mutation classes are *GC*-conservative, meaning that they are unaffected by gBGC. The other four mutation classes are *GC*-changing and include both transitions (*b*_*p*_, *c*_*p*_) and transversions (*d*_*p*_, *e*_*p*_). They are susceptible to be affected by gBGC; as stated before, transition and transversion mutations may be differently affected by gBGC (Lartillot, 2012; Bergman & Schierup, 2021). This justifies going beyond the usual classification of *GC*-changing (S↔ W) and *GC*-conservative (S ↔S or W↔ W) mutations (Bolívar et al., 2016; Boman et al., 2021).

We inferred the divergence with a *D*_*a*_-like measure for six nucleotide classes (*a*_*d*_, *b*_*d*_, *c*_*d*_, *d*_*d*_, *e*_*d*_, *f*_*d*_) as above (see equation 4). Since fluctuations in population size tend to converge to the harmonic mean over typical divergence times (Wright, 1940), for large populations, such as *Drosophila*, the influence of demography, *i*.*e*., changes in effective population size, on divergence is relatively small compared to the effects of directional forces like gBGC and selection (*e*.*g*., Kimura, 1962; Vogl & Mikula, 2021). As long as divergence times are relatively small, such that double mutations are too rare to influence inference, the independence of *a*_*p*_ and *f*_*p*_ polymorphisms from gBGC also extends to *A* : *T* and *C* : *G* divergence. Additionally, when divergence times are large enough, little shared heterozygosity is expected. With these conditions met, divergence and polymorphism ratios can be compared between nucleotide classes and chromosomal regions to disentangle the effect of population genetic forces.

Generally, the expected shared heterozygosity *H*_*s*_ between two populations decreases at a rate of *t/N* (*i*.*e*., *H*_*s*_*e*^−*t/N*^), such that after *t* = *N* generations, the proportion of shared polymorphism would be 0.37. In the case of *D. melanogaster* and *D. simulans*, we estimate the *t/N* between about 4 − 19 (see table 2). Therefore, the proportion of heterozygosity shared between the species is expected to be between *e*^−4^ and *e*^−19^, *i*.*e*., between 0.02 and 0.00. Furthermore the probability of observing double mutations between two populations is approximately (2*μt*)^2^. An estimate of 2*μt* can be obtained by multiplying the estimated divergence *t/N* by the estimated expected heterozygosity *θ* = 4*μN* within a population. For *D. melanogaster*, the highest *θ* = 4*μN* is about 0.024, after excluding the peripheral regions with reduced diversity (table 1). Consequently, we expect the highest proportion of double mutations between *D. melanogaster* and *D. simulans* to be around (4.062 ·0.024*/*2)^2^ 0.002. We therefore treat shared polymorphism between the two species and double mutations as negligible and the twelve parameters (six mutation classes for scaled mutation rates and divergence, respectively) as independent. In the following, we will drop the subscripts when we refer to the mutation classes, as it is clear from the context whether polymorphism or divergence or ratios of both are implicated.

**Table 2:** Divergence over polymorphism ratios for *GC*-conservative (*a* and *f*) and *GC*-changing (*b, c, d, e*) mutations for different parts of the autosomes and X-chromosome (95% CIs in brackets). WChr: Whole chromosome, CChr: Central regions of the chromosome arms, PChr: Peripheral regions of the chromosome arms.

|  | Autosome |  | X |  |
| --- | --- | --- | --- | --- |
|  | *GC*-conservative | *GC*-changing | *GC*-conservative | *GC*-changing |
| WChr | 5.104 (4.846, 5.394) | 8.774 (8.452, 9.095) | 6.191 (5.280, 7.263) | 9.246 (8.410, 10.195) |
| CChr | 4.062 (3.819, 4.334) | 6.854 (6.565, 7.130) | 5.534 (4.715, 6.502) | 8.201 (7.353, 9.184) |
| PChr | 10.908 (9.659, 12.288) | 19.065 (17.570, 20.414) | 13.142 (8.606, 21.736) | 17.167 (13.276, 21.695) |

### Diversity estimates along chromosomes

Our estimates of the overall expected heterozygosity (see equation 81 Vogl et al., 2020) at 5SI sites without differentiating the nucleotides are identical to the results of previous studies (Parsch et al., 2010; Jackson & Charlesworth, 2021): X chromosomal heterozygosity is slightly higher or equal to the autosomal heterozygosity and for all chromosomes estimates are lower towards the peripheral regions (telomeres and centromeres) of chromosome arms (table 1). The latter is expected as in the *Drosophila* genome recombination rates are lower towards telomeres and centromeres (Comeron et al., 2012) and nucleotide polymorphism is reduced in regions with reduced recombination (Begun & Aquadro, 1992). It is also known that mutations are *AT* biased in *Drosophila* (Vogl & Bergman, 2015) and accordingly our estimates of mutation rates going from *G* or *C* to *A* or *T* (*b, d*) are higher than in their reverse direction (*c, e*), respectively, *i*.*e*., *b > c* and *d > e* (figure 2). The degree of this bias is lower for the X chromosome (*β*_*A*_ = 0.668 (95% CI: 0.664 − 0.671) and *β*_*X*_ = 0.621 (95% CI: 0.613− 0.630)). Additionally, estimates on average are higher for transitions (*b, c*) than for transversions (*a, f, d, e*) with a ratio of 2.18 (95% CI: 2.09 − 2.27). Note that the estimate of *b*, both a *GC*-to-*AT* mutation and a transition, is the highest.

**Figure 2:**
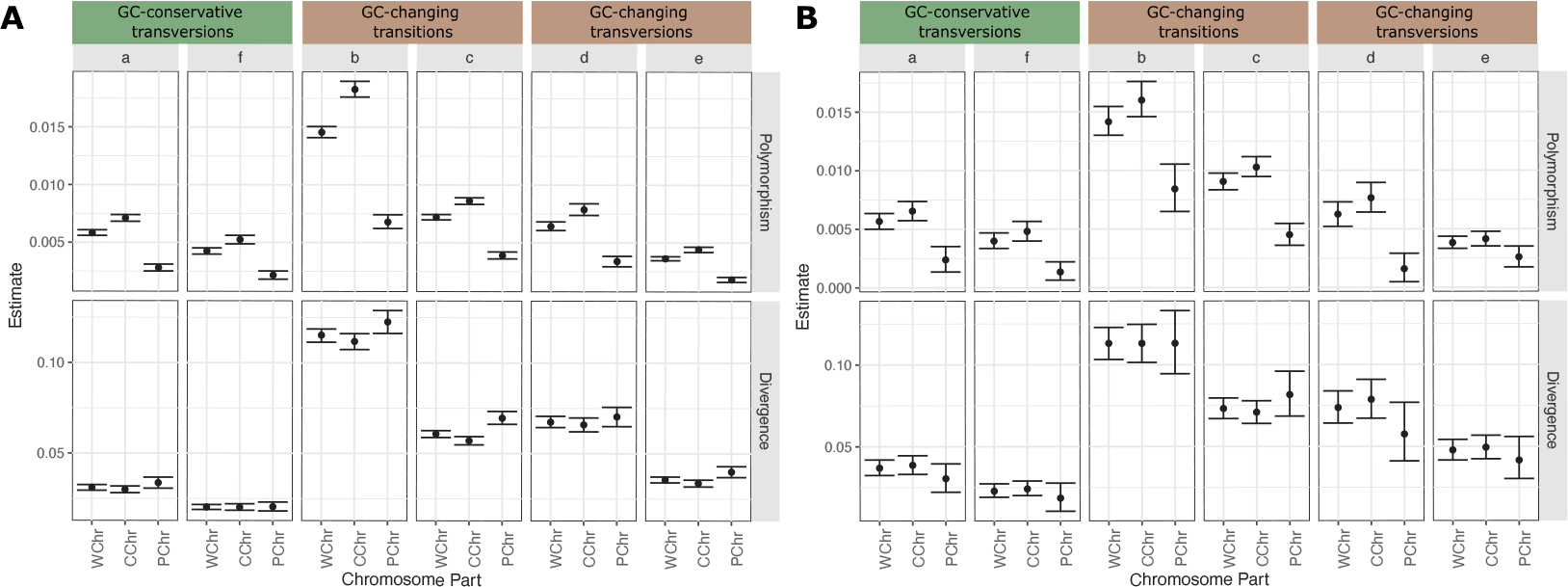
Scaled mutation rate (polymorphism) and divergence estimates of 5SI for the six mutation classes from the different parts of A) autosomes and B) X chromosome, where *a* and *f* are *GC*-conservative, *b* and *c GC*-changing transitions and *d* and *e GC*-changing transversions. WChr: Whole chromosome, CChr: Central regions of the chromosome arms, PChr: Peripheral regions of the chromosome arms.

The relative relationship between the nucleotide classes for the divergence estimates mirrors the patterns in scaled mutation rates. However, contrary to scaled mutation rates, divergence does not decrease towards the periphery, instead there is a slight increase for some *GC*-changing mutations in autosomes. This shows that mutation rate variation does not greatly contribute to a positive correlation between recombination and polymorphism, rather the variation in the effective population size *N*_*e*_ due to the direct or indirect effect of directional forces must cause it. If the major driver of the reduction in polymorphism levels is linked selection, we do not expect differences between mutation classes, while a directional force, like *GC*-biased gene conversion (gBGC), might differentially affect mutation classes. The presence of gBGC should be apparent from the differences between *GC*-changing (*b, c, d, e*) and *GC*-conservative (*a, f*) mutations. Indeed, we observe such differences, primarily driven by the *GC*-changing mutation class *b*. We also note differences between *GC*-conservative and other *GC*-changing mutation classes, although to a lower extent. However, comparing directly polymorphism or divergence estimates between mutation classes is not helpful: as *GC*-changing mutations include both transitions and transversions but *GC*-conservative ones only transversions, we still would not be able to distinguish between the effects of mutational and directional (gBGC) forces. Thus, for each mutation class, we get the ratio of divergence over polymorphism, which is minimally affected by mutation and is expected to scale inversely with the effective population size.

For all chromosomes and chromosomal parts, the divergence over polymorphism ratio of *GC*-changing mutations is significantly higher compared to *GC*-conservative ones (table 2). This shows that linked selection should not be the only driver creating nucleotide polymorphism variation along the genome and supports the presence of a directional force differing between these two classes of mutations, likely gBGC. But a directional force differing between them should result in a lower divergence to polymorphism ratio in classes with a directional force in equilibrium. We attribute this deviation from the equilibrium prediction (see Materials and Methods) to demography, in particular to recent population growth (Johri et al., 2020). Furthermore, the ratios do not differ significantly between *GC*-conservative mutations (*a, f*), yet among the *GC*-changing mutations they are slightly higher for transversions (*d, e*) than for transitions (*b, c*) (figure S1). This might be due to a difference in the strength of gBGC between transitions and transversions, which has been shown to be the case in other organisms (Lartillot, 2012; Bergman & Schierup, 2021).

In summary, our analysis of nucleotide polymorphism and divergence patterns along chromosomes show that linked selection can only account for part of the pattern. Varying divergence over polymorphism ratios in different mutation classes indicate an additional force, likely the non-adaptive directional force of gBGC.

### Diversity estimates between autosomes and the X chromosome

Under neutral equilibrium conditions (*e*.*g*., no mutational or effective population size difference between sexes) the expected divergence ratio of X over autosomes should be one and the expected polymorphism ratio 3*/*4 = 0.75. However, previously reported estimates of the X/A neutral site diversity in the ancestral populations of *Drosophila melanogaster* are approximately one (Campos et al., 2013). This was explained by linked selection, specifically by background selection (BGS): a higher recombination rate in the X chromosome should counteract the effect of BGS leading to less reduction in diversity. To support this argument, it was reported that the observed ratio of X/A diversities are recovered when BGS is modelled with the estimates of the distribution of fitness effect of deleterious mutations (Charlesworth, 2012; Comeron, 2014). Furthermore, when regions with similar effective recombination rates are compared, the ratio of mean X/A diversity values was shown to be close to the expected value of 0.75 (Vicoso & Charlesworth, 2009b; Campos et al., 2013, 2014). However, our analyses above showed that mutation classes might be affected distinctly by different forces, thus we investigated the X/A ratios for polymorphism and divergence separately for each mutation class.

Except for the peripheral regions, both polymorphism and divergence ratios are generally higher than their expected values (0.75 and 1, respectively, table 3). Yet, the deviation in X/A polymorphism ratios can not be only driven by BGS as suggested before, both because there are differences between mutation classes and the mutation rates are also slightly higher for the X chromosome compared to autosomes. Can these deviations be explained by mutation rate differences between X and autosomes? In the case of a pure mutation rate effect, polymorphism and divergence estimates of the chromosomes increase or decrease proportionally, so that the ratio between these two estimates should be unaffected. Thus without the effect of any directional force, we would expect *X*_*pol*_*/A*_*pol*_ = 0.75 *X*_*div*_*/A*_*div*_ (See also equations 5 and b).

**Table 3:**
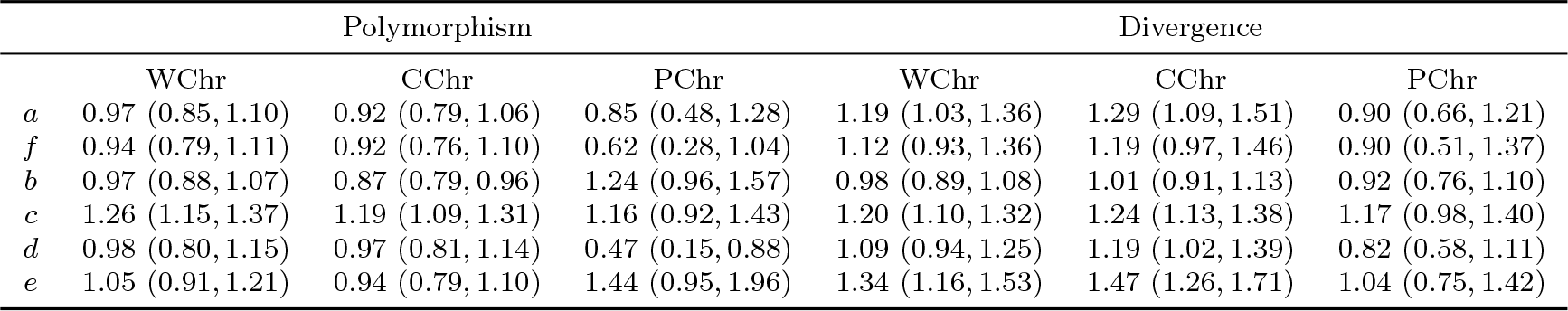
X/A ratios for polymorphism and divergence estimates of 5SI for each mutation class from the different parts of chromosomes (95% CIs in brackets); *a* and *f* are *GC*-conservative, *b* and *c GC*-changing transitions and *d* and *e GC*-changing transversions; WChr: Whole chromosome, CChr: Central regions of the chromosome arms, PChr: Peripheral regions of the chromosome arms.

|  | Polymorphism |  |  | Divergence |  |  |
| --- | --- | --- | --- | --- | --- | --- |
|  | WChr | CChr | PChr | WChr | CChr | PChr |
| *a* | 0.97 (0.85, 1.10) | 0.92 (0.79, 1.06) | 0.85 (0.48, 1.28) | 1.19 (1.03, 1.36) | 1.29 (1.09, 1.51) | 0.90 (0.66, 1.21) |
| *f* | 0.94 (0.79, 1.11) | 0.92 (0.76, 1.10) | 0.62 (0.28, 1.04) | 1.12 (0.93, 1.36) | 1.19 (0.97, 1.46) | 0.90 (0.51, 1.37) |
| *b* | 0.97 (0.88, 1.07) | 0.87 (0.79, 0.96) | 1.24 (0.96, 1.57) | 0.98 (0.89, 1.08) | 1.01 (0.91, 1.13) | 0.92 (0.76, 1.10) |
| *c* | 1.26 (1.15, 1.37) | 1.19 (1.09, 1.31) | 1.16 (0.92, 1.43) | 1.20 (1.10, 1.32) | 1.24 (1.13, 1.38) | 1.17 (0.98, 1.40) |
| *d* | 0.98 (0.80, 1.15) | 0.97 (0.81, 1.14) | 0.47 (0.15, 0.88) | 1.09 (0.94, 1.25) | 1.19 (1.02, 1.39) | 0.82 (0.58, 1.11) |
| *e* | 1.05 (0.91, 1.21) | 0.94 (0.79, 1.10) | 1.44 (0.95, 1.96) | 1.34 (1.16, 1.53) | 1.47 (1.26, 1.71) | 1.04 (0.75, 1.42) |

We plot X/A ratios for polymorphism and divergence, after adjusting divergence ratios by multiplying them with 0.75 to account for the expectation of *X*_*pol*_*/A*_*pol*_ = 0.75 *X*_*div*_*/A*_*div*_. The overlap between the X/A ratios for polymorphism and for adjusted divergence confirms that this expectation holds for *GC*-conservative mutations (*a, f*) (figures 3, S2), which seem to evolve under purely neutral forces and the higher X/A polymorphism ratio can be explained by a high X-chromosomal mutation rate in these nucleotide classes without invoking the effect of linked selection. Compared to autosomes, the X chromosome has relatively higher *AT* -to-*GC* mutation rates (*c, e*). This finding is unsurprising, as we have previously reported a reduced mutation bias towards *AT* on the X chromosome. However, nucleotide classes without mutation bias (*a, f*) also exhibit slightly elevated mutation rates in the X chromosome. This suggests that factors other than mutational bias contribute to the differences in mutation rates between chromosomes.

**Figure 3:**
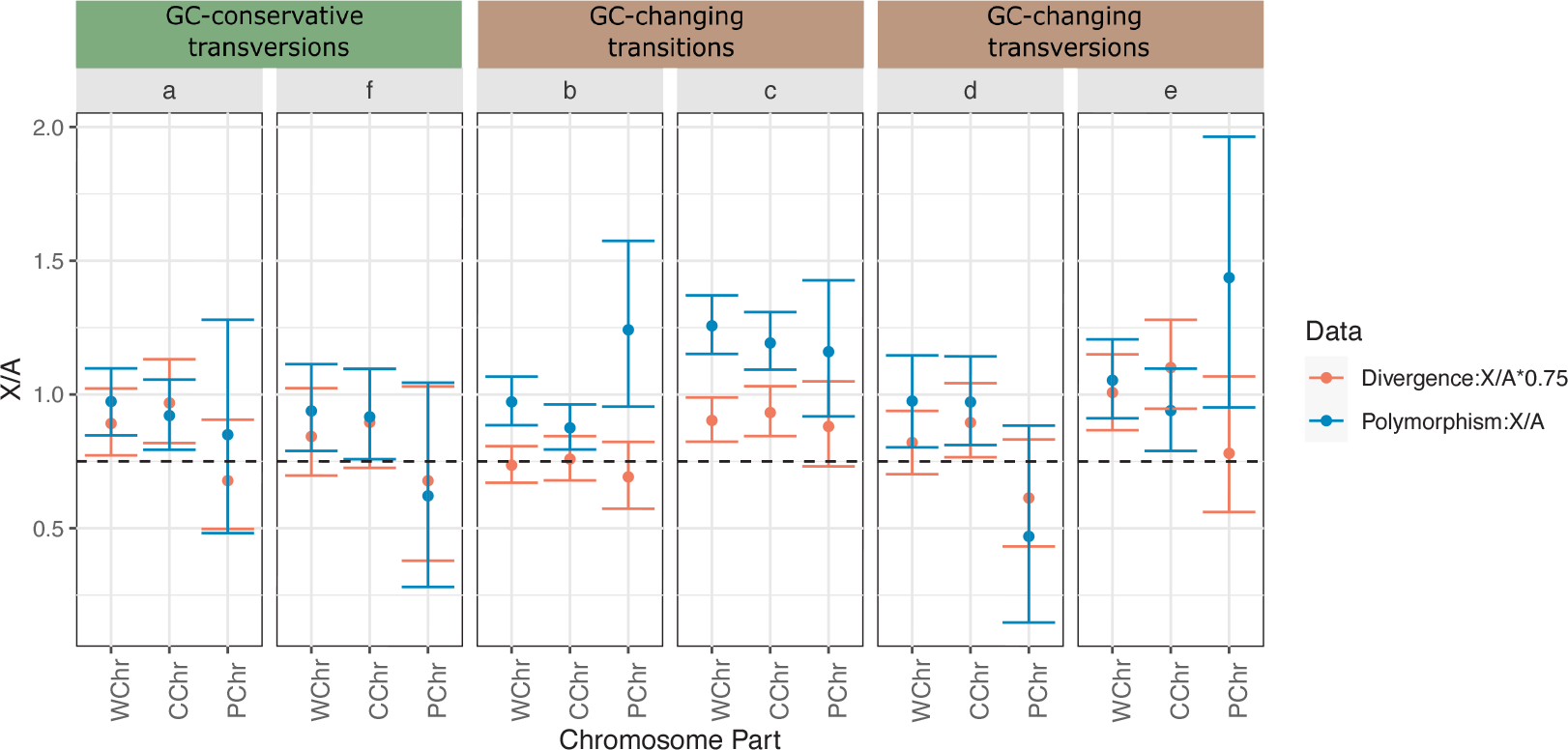
X/A ratios for polymorphism and divergence estimates. The divergence ratios are multiplied with 0.75 to account for the expectation of *X*_*pol*_*/A*_*pol*_ = 0.75 *X*_*div*_ */A*_*div*_ among mutation classes; *a* and *f* are *GC*-conservative, *b* and *c GC*-changing transitions and *d* and *e GC*-changing transversions; horizontal dashed line correspond to the value of 0.75. WChr: Whole chromosome, CChr: Central regions of the chromosome arms, PChr: Peripheral regions of the chromosome arms.

While the *GC*-conservative mutation classes follow the neutral expectation *X*_*pol*_*/A*_*pol*_ = 0.75 *X*_*div*_*/A*_*div*_, the *GC*-changing mutation classes deviate from it (figures 3, S2). Nonoverlapping values between X/A ratios for polymorphism and adjusted divergence suggest that gBGC might influence the molecular evolution patterns of X and autosomes differently. The most extreme deviations are observed in mutation classes *b* and *c*, and to a lower extent, in class *e* at peripheral regions, indicating that differences between X and autosomes are primarily driven by gBGC acting on transitions and on telomeric and centromeric regions.

In summary, mutation rates differ between autosomes and the X chromosome. Upon accounting for these differences, *GC*-conservative mutations conform to neutral expectations. But the pattern of *GC*-changing mutations suggests gBGC in shaping chromosomal disparities. Thus, when comparing the evolution of autosomes and the X chromosome, analyses should either be restricted to *GC*-conservative mutation classes or gBGC needs to be accounted for.

### Diversity estimates in relation to recombination rates

Given the lack of variation in divergence levels, the variation in polymorphism levels along the genome has been explained by the effect of linked selection, thus with variation in *N*_*e*_, in *Drosophila* species (Begun & Aquadro, 1992). The effect of linked selection should not differ among mutation classes. Contrary to this expectation, we observed distinct patterns among mutation classes and along chromosomes that we attributed to gBGC (table 2). As both gBGC and linked selection should be tied to recombination, we next investigated the diversity estimates of six mutation classes for different recombination rates. For this, we combined introns with similar recombination rates and compared results among them. We created four bins with approximately equal numbers of observations and also performed the binning after excluding telomeres and centromeres. The mean recombination rates within bins are approximately equal for X and autosomes (table S1).

The relative relationship between the estimates of mutation classes follows the same patterns reported before: higher rates for transitions and *GC*-to-*AT* mutations (figure 4). Among the recombination bins including telomeres and centromeres, polymorphism estimates decrease with decreasing recombination rate, which is much more pronounced for autosomes. However, the varia-tion in divergence between recombination bins is not significant, showing once again that mutation rate variation does not significantly contribute to the positive correlation between polymorphism and recombination in *Drosophila*. When peripheral regions of the chromosome arms with very low recombination rates are excluded, the decrease in polymorphism levels diminishes relatively. This suggests that the forces contributing to the association between polymorphism and recombination are mainly caused by differences between the central and peripheral regions of the chromosome arms.

**Figure 4:**
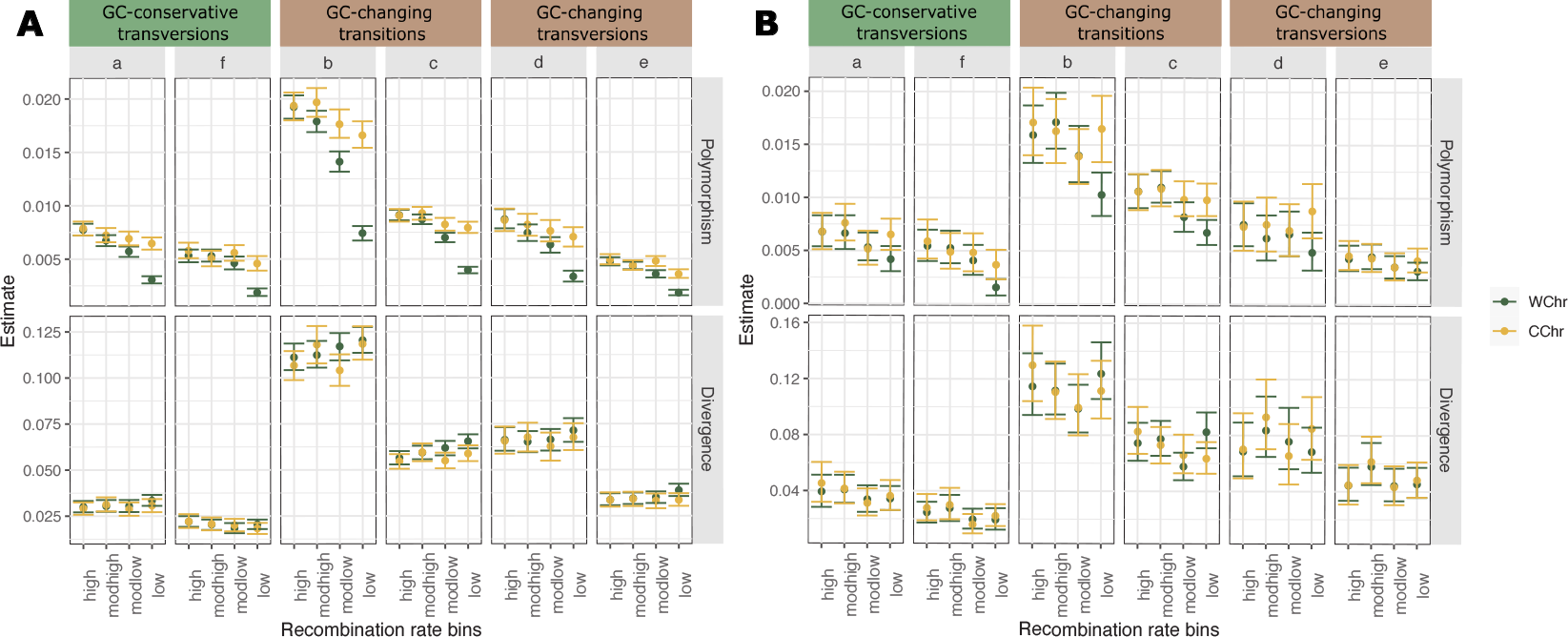
Scaled mutation rate (polymorphism) and divergence estimates of 5SI for the six mutation classes from the different recombination rates of A) autosomes and B) X chromosome, where *a* and *f* are *GC*-conservative, *b* and *c GC*-changing transitions and *d* and *e GC*-changing transversions. Introns are binned by recombination rate before (green, WChr: Whole chromosome) and after excluding telomeres and centromeres (yellow, CChr: Central regions of the chromosome arms).

Next we asked to what extent the polymorphism-recombination relationship is caused by linked selection and possibly by gBGC. We factored out mutational differences by calculating the ratio of divergence over polymorphism. In whole chromosomes we find that the lowest recombination rate class has significantly higher ratios than all other classes (figure 5). Comparing *GC*-conservative (*a, f*) and changing (*b, c, d, e*) mutations, we note that this effect is strongest in *GC*-changing mutations. Thus gBGC seems to contribute to the positive correlation between nucleotide polymorphism and local rates of recombination. The other recombination classes differ little from each other. After excluding telomeres and centromeres, variation is only significant for the lowest recombination bin of *GC*-changing mutations and surprisingly, the difference between *GC*-changing transitions and transversions is also lost. In contrast, no differences among recombination classes can be found for *GC*-conservative mutations.

**Figure 5:**
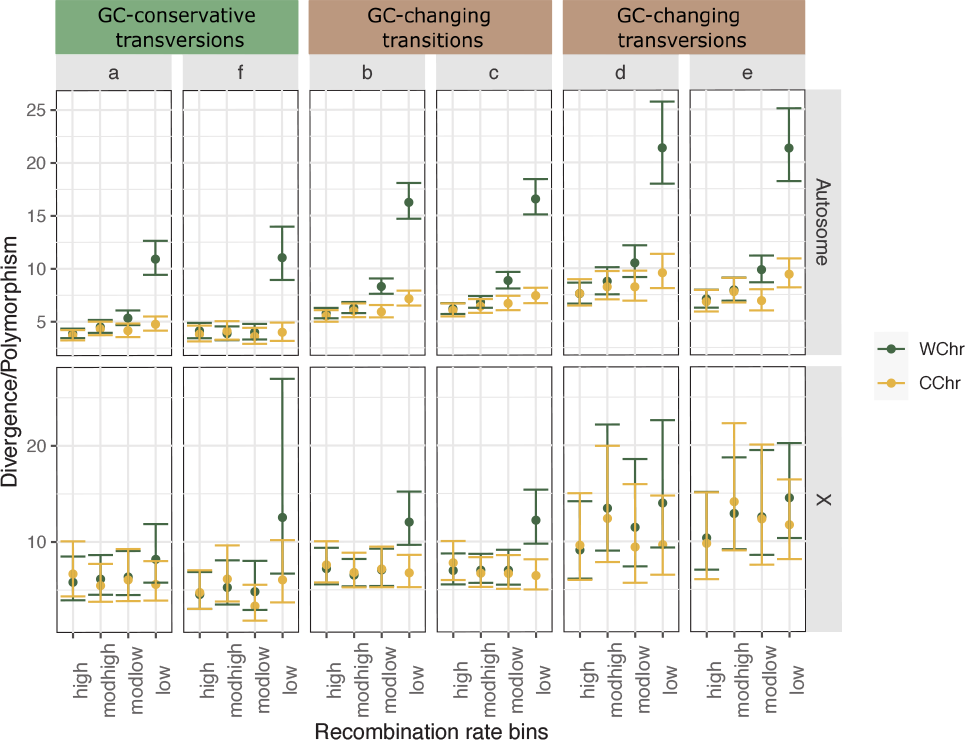
Divergence over polymorphism ratios for all six mutations classes from autosomes and X-chromosome, where *a* and *f* are *GC*-conservative, *b* and *c GC*-changing transitions and *d* and *e GC*-changing transversions. Introns are binned by recombination rate before (green, WChr: Whole chromosome) and after excluding telomeres and centromeres (yellow, CChr: Central regions of the chromosome arms).

In summary, divergence over polymorphism ratios differ among recombination classes only when the peripheral regions of chromosome arms are included. Variation among recombination classes within the central regions is low. Particularly, there is no trend toward increasing ratios (and thus presumably to decreasing effective population sizes due to linked selection) across different recombination rate classes within the central chromosome arms in *GC*-conservative mutation classes, while such a trend is discernible but barely significant in *GC*-changing mutation classes.

Lastly, we checked if the X/A diversity ratios among recombination classes conform to the expectation of *X*_*pol*_*/A*_*pol*_ = 0.75 *X*_*div*_*/A*_*div*_. As above, polymorphism ratios are generally higher than or equal to 3/4 and the patterns change between *GC*-changing and conservative classes (figure 6). For *GC*-conservative mutations, the observed ratio is totally explained by the mutation rate differences when telomeres and centromeres are excluded and deviates only slightly for the lowest recombination bin when they are included. It seems that the effect of background selection is creating a difference in X and autosome variation patterns only in centromeres/telomeres. For *GC*-changing mutations, there are again deviations from the neutral expectation, stronger towards low recombination rates and for transitions (figure S3). When telomeres and centromeres are excluded, the deviation is still observed for the lowest recombination bin (figure 6) due to transitions (figure S4). These results are in line with the findings above: the effect of gBGC on X and autosome differs mostly for transitions and in very low recombining regions.

**Figure 6:**
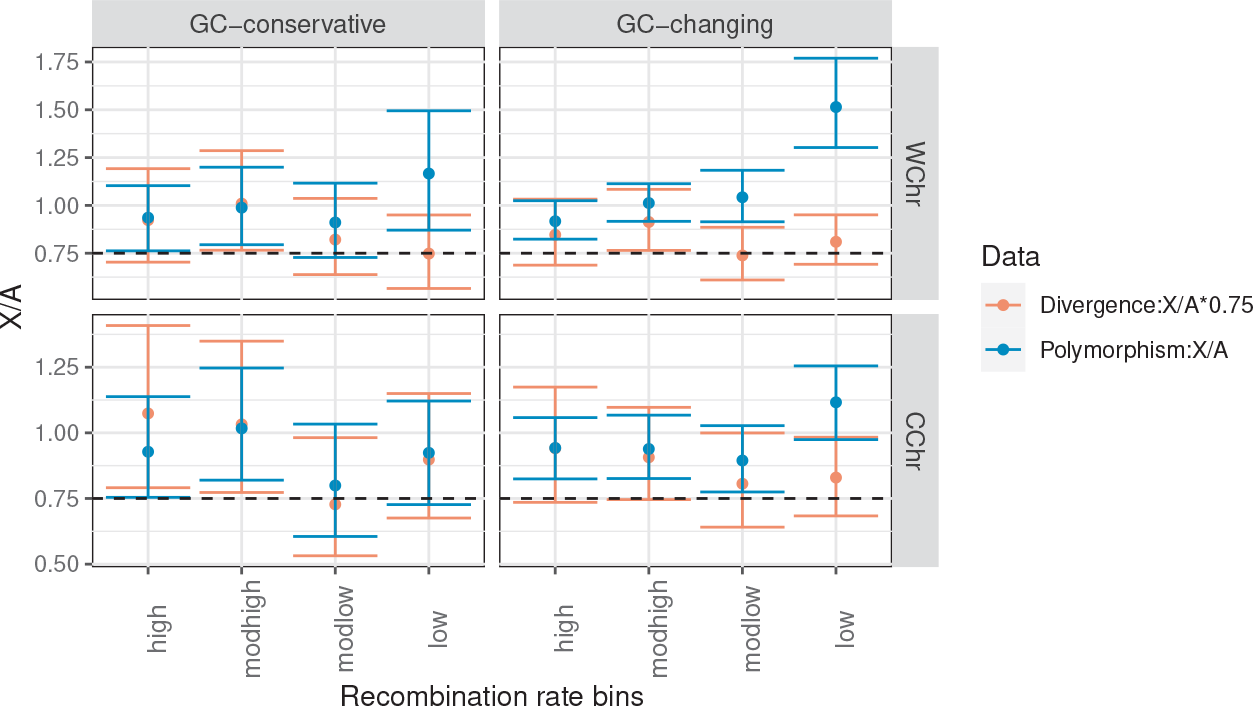
X/A ratios for polymorphism (blue) and divergence (red) estimates of *GC*-conservative (*a* and *f*) and changing (*b, c, d, e*) mutations. Introns are binned by recombination rate before excluding telomeres and centromeres (top panel, WChr: Whole chromosome) or after excluding telomeres and centromeres (lower panel, CChr: Central regions of the chromosome arms). The divergence ratios are multiplied with 0.75 to account for the expectation of *X*_*pol*_*/A*_*pol*_ = 0.75 *X*_*div*_ */A*_*div*_ among mutation classes. The horizontal dashed line correspond to the value of 0.75.

Previous studies showed that the X/A diversity ratio is close to the neutral expectation when regions with equal effective recombination rates are compared (*i*.*e*., the rates for the X chromosome are 4*/*3 of the rates for autosomes due to lack of recombination in male *Drosophila*) with an exception of regions with very low recombination (Vicoso & Charlesworth, 2009b; Campos et al., 2013, 2014). They suggested that this further supports BGS as the main driver of the high X/A diversity ratio when whole chromosomes are considered and as long as regions with similar recombination rates are compared, expectations for neutral diversity should be met. However, we observed that patterns differ between *GC*-conservative and *GC*-changing mutations. Specifically, after accounting for mutational differences, there is no deviation from neutral equilibrium expectations in the *GC*-conservative mutations in the central regions of chromosome arms. Therefore BGS appears not to be the driver of chromosomal differences, except at telomeres and centromeres, while the effect of gBGC should be considered.

### Strength of gBGC

As we find evidence for the effect of gBGC in diversity patterns, we next inferred its strength, quantified as *B* = 4*N*_*e*_*b*, using the ML estimator of Vogl & Bergman (2015). We obtained estimates either jointly for all *GC*-changing mutations (*B*_*GC*_) or separately for *GC*-changing transitions (*B*_*Ts*_) and transversions (*B*_*Tv*_) from SFS constructed based on segregating *GC*-frequency. We tested whether separately considering transitions and transversions improved the fit by a likelihood ratio test, where we compared the likelihood of *B*_*GC*_ to the sum of the likelihoods of *B*_*Tv*_ and *B*_*Ts*_. The difference between the estimates of autosomes and the X chromosome or between the peripheral and central parts of chromosomes are evaluated in a similar manner.

The force favoring *GC* is weak, *i*.*e*., about *B* ≈ 0.5, but differs significantly from zero both in autosomes and the X chromosome (table 4). Estimates are also significantly different from zero if transitions and transversions are analyzed separately (table 5). For central autosomal arms, *B*_*Tv*_ is significantly stronger than 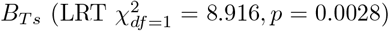, while for peripheral regions, the difference is not significant 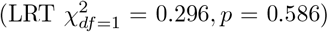. *B*_*Tv*_ estimates do not differ between the central and peripheral parts of the chromosome arms 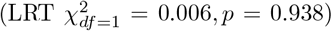, while *B*_*Ts*_ estimates do 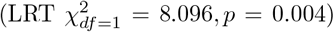. Conversely, for the X chromosome, *B*_*Ts*_ and *B*_*Tv*_ differ significantly at the peripheral region of the chromosome arms, with a higher 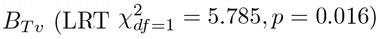, while at the central part the value for all *GC*-changing mutations fits the data better 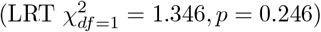. This is explained by an increasing *B*_*Tv*_ value towards telomeres and centromeres, as *B*_*Ts*_ does not change significantly along the X chromosome 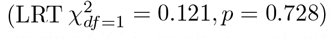, while *B*_*Tv*_ does 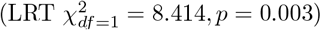.

**Table 4:** *B* values inferred from the SFS of *GC*-changing mutations (*B*_*GC*_), for autosomes, the X chromosome, and autosomes and X pooled (95% CIs constructed from likelihood ratio test are given in brackets). Estimates are either given separately for central and peripheral regions of the chromosome arms (CChr and PChr, respectively) or for whole chromosomes (WChr). All values are significantly different from *B* = 0 (*p <* 0.001). Bold values for the pooled data indicate no significant differences between autosomes and the X.

|  | WChr | CChr | PChr |
| --- | --- | --- | --- |
| Autosomes | 0.504 (0.451, 0.558) | 0.477 (0.419, 0.535) | 0.650 (0.515, 0.787) |
| X | 0.597 (0.451, 0.743) | 0.564 (0.410, 0.719) | 0.853 (0.422, 1.30) |
| Pool | **0.515 (0.465, 0.565)** | **0.488 (0.434, 0.542)** | **0.668 (0.539, 0.799)** |

**Table 5:** *B* values inferred from the SFS of *GC*-changing transitions (*B*_*Ts*_) and transversions (*B*_*Tv*_), for autosomes, the X chromosome, and autosomes and the X pooled (95% CIs constructed from likelihood ratio test are given in brackets). Estimates are either given separately for central and peripheral regions of the chromosome arms (CChr and PChr, respectively) or for whole chromosomes (WChr). All values are significantly different from *B* = 0 (*p <* 0.001). Bold values for the pooled data indicate no significant differences between autosomes and the X.

|  | WChr |  | CChr |  | PChr |  |
| --- | --- | --- | --- | --- | --- | --- |
|  | Transitions | Transversions | Transitions | Transversions | Transitions | Transversions |
| Autosome | 0.457 (0.392, 0.522) | 0.605 (0.510, 0.699) | 0.417 (0.346, 0.487) | 0.606 (0.503, 0.710) | 0.677 (0.512, 0.843) | 0.596 (0.359, 0.835) |
| X | 0.613 (0.439, 0.789) | 0.559 (0.295, 0.826) | 0.625 (0.439, 0.812) | 0.425 (0.148, 0.706) | 0.529 (0.029, 1.04) | 1.781 (0.894, 2.78) |
| Pool | 0.476 (0.415, 0.537) | **0.600 (0.511, 0.689)** | 0.443 (0.377, 0.509) | **0.585 (0.488, 0.682)** | **0.662 (0.506, 0.821)** | 0.682 (0.454, 0.913) |

On both autosomes and the X chromosome, there is an increase in the overall strength of gBGC (*B*_*GC*_) towards the telomeres and centromeres (table 4). This is due to a significant increase in *B*_*Ts*_ for autosomes, while *B*_*Tv*_ shows a significant increase for the X chromosome. To assess whether these changes along the chromosomes cause significant differences between the X chromosomal and autosomal estimates of gBGC, we compared pooled data to their estimates via LRT. For the central part of the chromosome arms, the difference between the X chromosome and autosomes is significant due to the differences in 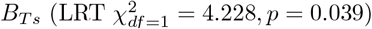, while for the peripheral regions, differences in *B*_*Tv*_ cause a significant deviation 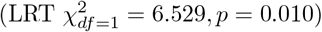. These results are consistent with those of diversity patterns, both in comparisons along chromosomes and between chromosomes.

We also estimated *B* values while accounting for the effect of demography through addition of correction parameters (*r*_*y*_) that are obtained from the neutral SFS, *i*.*e*., SFS of *GC*-conservative mutations (see Materials and Methods). We note that this decreases the statistical power. Nonetheless, all estimates remain significantly different from zero, and the values fall within a similar range as the estimates without correction (see Figure S5 and Tables S4 and S5). Furthermore, we still observe an increase of *B* towards telomeres and centromeres in *B*_*Ts*_ for autosomes and in *B*_*Tv*_ for X chromosomes, however, these trends are no longer statistically significant, likely due to the reduced power (figure S5).

That our estimates of *B* do not decrease near telomeres and centromeres compared to the central part of the chromosome arms, even after accounting for demographic disequilibrium, requires an explanation. *B* = 4*N*_*e*_*b* measures the strength of the directional force scaled by the effective population size. The conversion bias *b* depends on the average length of the conversion tract, the repair bias towards *GC*, and the recombination rate per site per generation. Notably, our recombination rate estimates rely on only crossover (CO) events. As CO rates and the effective population size decreases towards telomeres and centromeres (see the mutation classes *a* and *f* in the autosomes in figure 5), the conversion bias *b* needs to increase disproportionately to explain the patterns in *B* values. This suggests that *b* might be governed by a more intricate interplay of factors than a simple relationship with CO rates.

To show the relationship between gBGC and the recombination (CO) rate, we calculated *B* from the SFS data binned according to CO rates. Although there is a slight increase in *B* towards low recombining regions, when telomeres and centromeres are included, the estimates do not differ significantly among recombination bins (figure S6). Importantly, estimates inferred without recombination binning (table 4) provide a better fit to the data (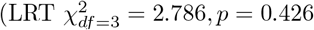 for autosomes; 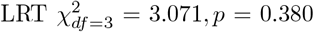 for the X chromosome). Furthermore, the *B* estimates obtained without assuming demographic equilibrium also do not exhibit any positive or linear relationship with CO rates. There is still a slight, non-significant increase in *B* towards low-recombining regions (figure S7). When telomeres and centromeres are included, estimates from the lowest recombination rate classes are similar to those from the highest recombination rates.

Our inference of gBGC strength so far relies on polymorphic spectra and is thus limited to currently segregating alleles. On the other hand, the general *GC* content including monomorphic sites is influenced also by earlier events that already fixed. Using the *GC* content thus promises more power for inference. A slight but significant negative correlation between *GC* content and CO rates is observed for the 5SI regions when analyzing whole chromosomes (Spearman’s *ρ* = − 0.048, *p <* 0.001 for autosomes; Spearman’s *ρ* = − 0.140, *p <* 0.001 for the X chromosome). After excluding telomeres and centromeres, the significance is lost for autosomes and reduced for the X chromosome (Spearman’s *ρ* = − 0.018, *p* = 0.192 for autosomes; Spearman’s *ρ* = − 0.116, *p* = 0.002 for the X chromosome).

While the CO rate and the strength of gBGC are expected to be indirectly related, the relationship between *GC* content and gBGC is expected to be direct. Consistent with our observation that gBGC is relatively constant along the chromosomes, the *GC* content of the *Drosophila* genome also shows little variation. Yet, when examining the differences in base composition, we may still expect a weak correlation between *B* values and *GC* content. In order to investigate this, it is necessary to group introns into bins according to their *GC* content. However, estimating the strength of gBGC from data binned according to its own *GC* content might create a bias. To avoid this dependence, we created five bins with approximately equal sizes by using the mean *GC* content of fourfold degenerate sites from the same gene where the short introns are located. This choice is supported by the significant and positive correlation between FFDS and 5SI *GC* content across all chromosomes (Spearman’s *ρ* = 0.257, *p <* 0.001, Spearman’s *ρ* = 0.273, *p <* 0.001 for autosomes and X chromosome, respectively), as also reported by previous studies (Kliman & Eyre-Walker, 1998; Galtier et al., 2006). Most of the data clustered in intermediate levels of *GC* content (figure S8), thus the *GC* content ranges of the bins were not equal (table S2). Furthermore the mean CO rates were similar among *GC*-bins (≈ 2.27 cM/Mb and ≈2.56 cM/Mb for autosomes and X chromosome, respectively). Even though a relationship between *GC* content, thus gBGC, and CO rates is expected, we once more fail to observe it.

The estimates of *B*_*Ts*_ and *B*_*Tv*_ do not significantly differ from each other, *i*.*e*., *B*_*GC*_ fits data better, for both chromosomes and for all *GC*-bins. More crucially, estimates from *GC*-bins explain our data better than the whole chromosome estimates (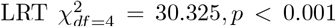 for autosomes; 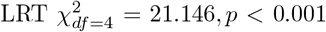 for X chromosome). Among the *GC*-bins, *B*_*GC*_ is higher for highest *GC* content, yet the difference among the other four classes is not significant (figure 7A). Due to the relatively uniform base composition along chromosomes, we expected little power to detect co-variation in *B* values, nevertheless we detected such an association. Additionally, while the strength of the directional force, *i*.*e*., gBGC, increases with *GC* content, the mutation bias towards *AT* is independent of it (figure 7B). Accounting for the demographic disequilibrium had no effect on the patterns between *GC*-bins, with the only impact being on the absolute values of *B* (figure S9).

**Figure 7:**
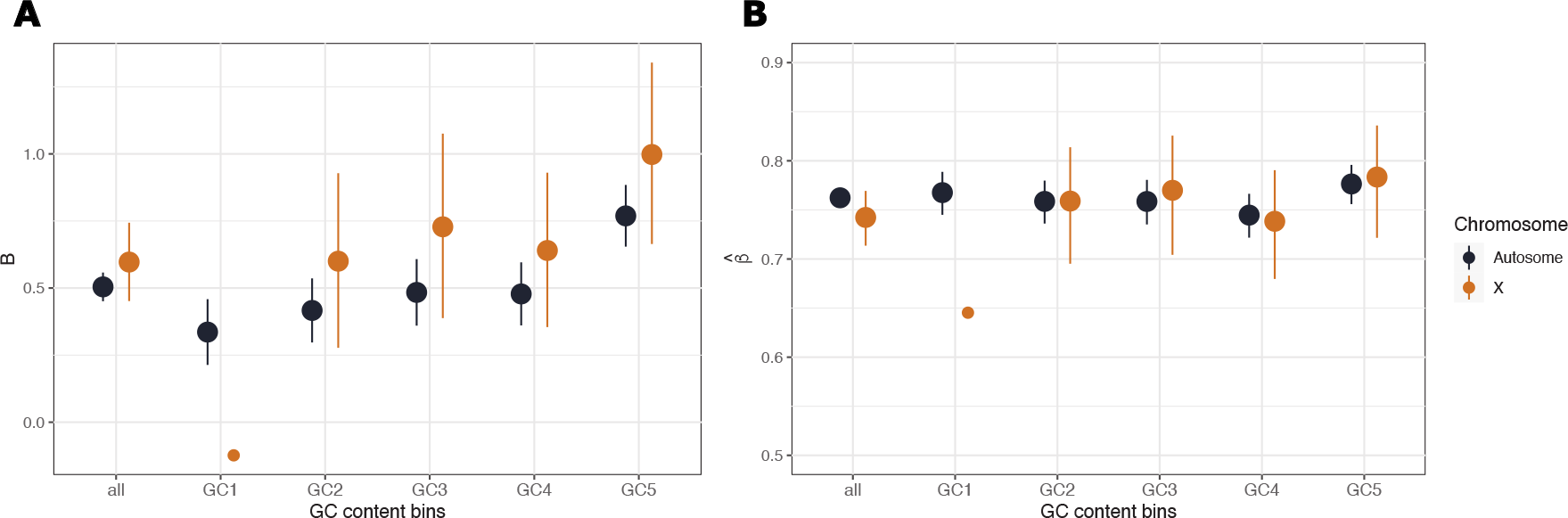
A) *B*_*GC*_ values inferred from the SFS of *GC*-changing mutations for autosomes (black) and X chromosome (brown). The big dots represents the estimates significantly different from *B* = 0. Confidence intervals are constructed from likelihood ratio test. B) Mutation bias estimated conditional on *B* for autosomes (black) and X chromosome (brown). Estimates are given for all introns and for introns binned by the mean *GC* content of the FFDS of the same genes. *GC* content increases from GC1 to GC5 and the ranges are given in S2

### Comparison to Drosophila simulans

So far we used polymorphism data from *D. melanogaster* and divergence data between *D. melanogaster* and *D. simulans*. As the shared polymorphism between the two species is negligible, we also analyzed the polymorphic spectrum of *D. simulans* to compare patterns of the different mutation classes between the species.

As in *D. melanogaster*, the inferred strength of gBGC is about *B* = 0.5 and significantly different from *B* = 0 (table 6). Estimates are higher for *GC*-changing transversions than for *GC*-changing transitions 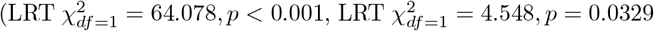 for autosomes and X, respectively). Importantly, divergence over polymorphism ratios are higher for *GC*-changing mutations than for *GC*-conservative ones and increase with the strength of gBGC, again as in *D. melanogaster* (figure S10).

**Table 6:** *B* values inferred from the SFS of all *GC*-changing mutations (*B*_*GC*_), *GC*-changing transitions (*B*_*Ts*_) and transversions (*B*_*Tv*_) for autosomes, the X chromosome, and pooled data (autosomes+X) of *D. simulans* population (95% CIs constructed from likelihood ratio test are given in brackets). All values are significantly different from *B* = 0 (*p <* 0.001). Bold values for the pooled data indicate no significant differences between autosomes and the X.

| | $B_{GC}$ | $B_{Tv}$ | $B_{Ts}$ |
| --- | --- | --- | --- |
| Autosomes | 0.459 (0.398, 0.520) | 0.763 (0.656, 0.869) | 0.311 (0.235, 0.387) |
| X | 0.345 (0.152, 0.524) | 0.588 (0.261, 0.921) | 0.232 (0.001, 0.464) |
| Pool | **0.447 (0.398, 0.496)** | **0.745 (0.659, 0.832)** | **0.303 (0.243, 0.362)** |

Furthermore, the *GC* content of four-fold degenerate and 5SI sites of the same gene are also correlated significantly and positively, as in *D. melanogaster* (Spearman’s *ρ* = 0.267, *p <* 0.001, Spearman’s *ρ* = 0.283, *p <* 0.001 for autosomes and the X chromosome, respectively). Thus, we also grouped the SFS into five equally sized bins depending on the background FFDS base composition and inferred *B* (table S3). The estimates are significantly greater than *B* = 0 for all *GC*-bins in autosomes and for the two bins with the highest *GC* content in the X chromosome (figure 8A). Among the significant estimates, the *B* values increase with *GC* content, while the mutation bias towards *AT* remains constant (figure 8B). Overall, patterns in *D. simulans* are similar to those in *D. melanogaster*. Once again, *B* values obtained after accounting for the disequilibrium follow the same patterns (table S6). However, the difference between *B*_*Tv*_ and *B*_*Ts*_ estimates and the estimates from the low *GC* bins on the X chromosome are not significant anymore (figure S11, S12).

**Figure 8:**
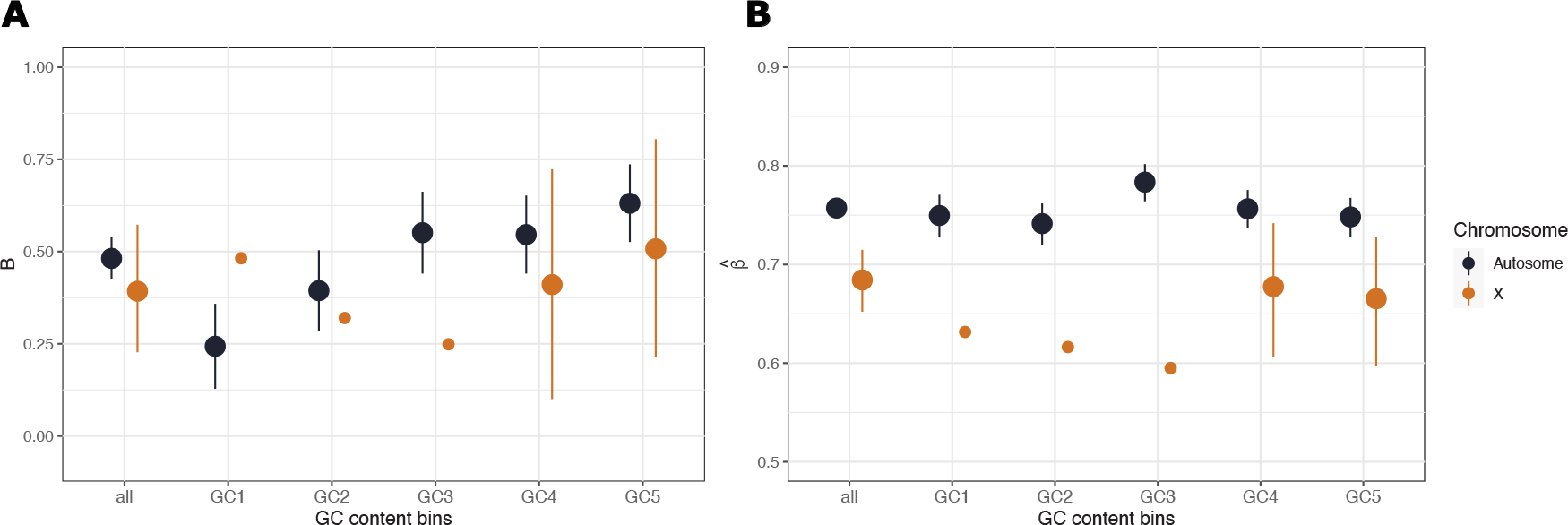
Results from *GC* content binned data of *D. simulans* population. A) *B*_*GC*_ values inferred from the SFS of *GC*-changing mutations for autosomes (black) and X chromosome (brown). The big dots represents the estimates significantly different from *B* = 0. Confidence intervals are constructed from likelihood ratio test. B) Mutation bias estimated conditional on *B* for for autosomes (black) and X chromosome (brown). Estimates are given for all introns and for introns binned by the mean *GC* content of the FFDS of the same genes. *GC* content increases from GC1 to GC5 and the ranges are given in S3

In summary, gBGC is present and affecting neutral sequence variation similarly in *Drosophila melanogaster* and *D. simulans*. This manifests in varying *GC* content along chromosomes, patterns of skewed site frequency spectra, and deviations in divergence to polymorphism ratios of *GC*-changing mutations compared to *GC*-neutral mutations.

## Discussion

Identifying the different nonadaptive evolutionary forces that act on putatively neutral sequences and describing their effects provides a better understanding of genome evolution patterns. These nonadaptive evolutionary forces can then be incorporated into null models to quantify their influence in the genome. By comparing the neutral patterns to genomic regions of functional importance, such models allow for the detection of adaptive forces. In such null models, the effect of linked selection has been included in addition to mutation and demography (Zeng & Charlesworth, 2010; Comeron, 2017; Johri et al., 2020), as it has been shown to shape patterns of genome variation in many organisms. More recently, the effect of *GC*-biased gene conversion (gBGC) has been demonstrated in many taxa (Pessia et al., 2012; Glémin et al., 2014; Galtier, 2021), including *Drosophila* (Jackson & Charlesworth, 2021). Failing to account for gBGC, when it is present, has been shown to lead to biased inference of selective and demographic forces (Bolívar et al., 2018; Pouyet et al., 2018), or lead to false interpretations about the effect of nonadaptive forces in shaping neutral sequence patterns (Bolívar et al., 2016; Boman et al., 2021). Thus, it is important to incorporate gBGC into the model when inferring adaptive forces and to use *GC*-conservative mutations for a more accurate representation of the effects of other nonadaptive factors governing neutral genome sequence evolution.

Short introns of *Drosophila*, specifically the 5’ sites of short introns (5SI), have been shown to evolve in the absence of selective constraints (Parsch et al., 2010; Halligan & Keightley, 2006). In *Drosophila*, most introns are short and evenly distributed across the chromosomes (Parsch et al., 2010), making them a good alternative to synonymous sites. Therefore 5SI sites have been increasingly used as a neutral reference to infer natural selection and population history (Lawrie et al., 2013; Garud et al., 2015; Machado et al., 2020), replacing synonymous sites, which can be influenced by codon usage bias (CUB) (Akashi, 1994). Nevertheless studies of 5SI sites have revealed evidence for a directional force, favouring the strong *GC* over the weak *AT* bases (Vogl & Bergman, 2015; Jackson et al., 2017). Understanding the cause of this pattern in neutral sequences is crucial both for comprehending the forces shaping genome evolution and ensuring accuracy of null models when assessing the impact of natural selection and population history. Recent research suggested that this *GC* preference might be due to gBGC (Jackson & Charlesworth, 2021), but it is still unknown how gBGC operates on different chromosomes or on different mutation classes (*e*.*g*., transitions vs transversions). Additionally, it is important to investigate how the presence of gBGC affects our prior interpretations regarding the effects of other nonadaptive forces on the evolution of neutral sequences in *Drosophila*.

Using the 5SI sites as neutrally evolving reference sequences in *Drosophila*, we find a pervasive influence of gBGC on patterns of neutral sequence variation in both *Drosophila melanogaster* and *D. simulans* that shows in variable *GC* content along chromosomes, correlated skewed polymorphism patterns, deviation of divergence to polymorphism ratios from predictions assuming only mutation and drift, and differences among transition and transversion mutations and between autosomes and the X chromosome. On the other hand, patterns in *GC*-conservative mutations show that predictions of the neutral theory are borne out, while many of the results formerly attributed to linked selection seem to actually be caused by gBGC.

Analysis of *GC*-changing mutation classes shows the presence of a directional force attributable to gBGC of about *B* ≈ 0.5. This is comparable to previous estimates from noncoding regions (Galtier et al., 2006) and short autosomal introns of *Drosophila* (Jackson & Charlesworth, 2021). Going beyond these earlier studies, we reveal that gBGC operates in all chromosomal regions of autosomes and the X chromosome and in *GC*-changing transitions and transversions, albeit with slightly varying strength among regions. Between autosomes and the X chromosome the pattern is complicated: transitions in the central regions and transversions in the peripheral regions are higher in the X chromosome (table 5), yet after accounting for deviations from neutral equilibrium, differences are not significant (figure S5). Along chromosomes, the strength of gBGC slightly increases towards telomeres and centromeres, where crossover (CO) rates are low (Comeron et al., 2012). This relationship between CO rates and the strength of gBGC is also reflected in the negative relationship between 5SI *GC* content and CO rates.

In *Drosophila*, noncrossover (NCO) and crossover (CO) rates are negatively correlated (Langley et al., 2000; Comeron et al., 2012) and NCO rates exhibit a more uniform distribution along the chromosomes compared to CO rates (Comeron et al., 2012; Miller et al., 2016). Our data indicate a higher or similar values of directional force *B* = 4*N*_*e*_*b* towards peripheral regions of chromosome arms where CO rates and effective population sizes *N*_*e*_ are low (*N*_*e*_ is here estimated independently from the *GC*-conservative mutations). This combination of observations suggests that the varying strength of the conversion bias *b* is not only associated with COs but also with NCOs in *Drosophila*. Such an association would explain both the negative relationship between 5SI *GC* content and CO rate and the relatively uniform strength of gBGC (and as a corollary the relatively uniform *GC* content in 5SI), except between the central and peripheral part of the chromosome arms. Previous studies failed to find a negative association between the *GC* content of introns and the CO rate in *D. melanogaster*, but rather reported weak positive correlations when considering whole genome *GC* content (Marais et al., 2003; Singh et al., 2005a). This might be due to an incomplete annotation of the reference genome, due to data excluding telomeres and centromeres, or because sites affected by directional selection were included. To our knowledge, only two studies reported a negative correlation between non-coding *GC* content and recombination in the X chromosome of *D. melanogaster* (Singh et al., 2005b; Campos et al., 2013). They also suggested higher gBGC in regions of low recombination as a possible explanation among others, however, discarded this possibility due to the apparent absence of such a relationship on the autosomes. As we observe such a negative relationship between crossover rate and *GC* content on both autosomes and the X chromosome, however, we uphold this hypothesis.

Some of the patterns of diversity and skew in the site frequency spectra observed in our study could also be explained by a recent change in the mutation bias instead of gBGC. We next summarize the arguments for gBGC using our data and analyses: while the genome of *D. melanogaster* has become more *AT* -rich compared to the ancestral state (Kern & Begun, 2005; Jackson & Charlesworth, 2021), which may explain a *GC*-skewed polymorphic SFS via a change in mutation bias, that of *D. simulans* has not (Jackson & Charlesworth, 2021), but nevertheless shows similarly skewed polymorphic SFS and diversity patterns as *D. melanogaster* (table 6, figures 8, S10). Furthermore, both in *D. melanogaster* and in *D. simulans* the *GC* proportion varies over the genome. The *GC* content of introns is correlated along chromosomes with that of fourfold degenerate sites (FFDS), pointing to a common mechanism. Using the polymorphic site frequency spectra (SFS), we inferred gBGC of varying strength in the 5SI correlated with *GC* content (figures 7, 8, S9, S12), but a rather uniform mutation bias that cannot explain variation in *GC* proportion. In addition, unlike Jackson & Charlesworth (2021), who partitioned their data based on the *GC* content of 5SI, which may introduce bias, we instead utilized the *GC* content of FFDS. Our observation of stronger *B* values for higher *GC* content provides further support for the common mechanism being gBGC. Thus, our evidence strongly points to gBGC as the more plausible explanation over a change in mutation bias.

Although the directional force of gBGC is within the nearly neutral range (Tachida, 1991), it has an impact on diversity patterns and can lead to false interpretations of genome evolution if not properly accounted for. *GC*-conservative mutation classes are not affected by gBGC and therefore suited to infer the effects of linked selection. We infer lower effective population sizes towards telomeres and centromeres than in central regions. This correlates with the overall pattern of CO rates. On the other other hand, variation in the local effective population size *N*_*e*_ in the central chromosomal regions is small and not significantly correlated with CO rates (figure 5). These findings replicate earlier showing the significant impact of recombination through linked selection, in generating variation between central and peripheral regions (Begun & Aquadro, 1992; Comeron et al., 2012). However, in contrast to these earlier studies we show little effect of linked selection on the variation in diversity patterns in central chromosomal regions (*e*.*g*., Cutter & Payseur, 2013; Comeron, 2014; Elyashiv et al., 2016). This has important consequences for the current efforts building null models: Within the central regions of chromosomes the CO map has little predictive value explaining the observed sequence patterns.

Comparing autosomes and the X chromosomes in *D. melanogaster* using *GC*-conservative mutations shows that, while the overall mutation rate is higher on the X, the effective population size of the X is about 3*/*4th that of the autosomes in the central region of chromosome arms (table 3 and figures 3,6, S2). The exact biological mechanism for the higher X mutation rate is unclear. Higher female mutation rate or differences in heterogametic male X chromosome, like dosage compensation (Gupta et al., 2006; Lucchesi & Kuroda, 2015) and distinct repair properties, might cause increase in mutation rates. Previous studies attributed differences between autosomes and X to background selection rather than mutation (Vicoso & Charlesworth, 2009a; Charlesworth, 2012; Campos et al., 2013; Comeron, 2014). We note that background selection does not create chromosomal differences in the central chromosomal regions, but only in the peripheral regions with very low recombination and is not the major driver of the patterns. Our finding is important for studies comparing evolutionary patterns between the X chromosome and autosomes: in the central regions of chromosome arms, the X/A ratio of neutral diversity in *GC*-neutral mutation classes is just as expected from the neutral theory, after accounting for differences in mutation rates. Failing to differentiate among mutation classes might obscure this simple pattern.

Differentiating between mutation classes allows us to tease apart the influence of various population genetic forces relevant for neutral sequence evolution patterns in *Drosophila*: the influence of mutation rates in different mutation classes and their variation, the influence of differences in recombination rates and linked selection, and the influence of gBGC. None of these individual forces dominates, rather they act jointly and interdependently. In our study, we took the joint influence of all weak forces on neutral sequence evolution in *Drosophila* into account. We believe that doing so will also improve the study of neutral and nearly neutral sequence evolution in other species.

## Acknowledgements

The authors wish to thank all members of the Vienna Graduate School of Population Genetics for support, and Lynette Caitlin Mikula for critically reading the article and providing helpful suggestions. We also thank two anonymous reviewers for their helpful comments on the manuscript. This work was supported by the Austrian Science Fund (FWF; W1225-B20).

## 1. Supplement

**Table S1:** Mean recombination rates (in cM/Mb) of datasets binned before and after excluding telomeres and centromeres. WChr: Whole chromosome, CChr: Central regions of the chromosome arms

|  | Autosome |  | X |  |
| --- | --- | --- | --- | --- |
|  | WChr | CChr | WChr | CChr |
| High | 4.89 | 5.49 | 5.33 | 5.78 |
| Modhigh | 2.54 | 3.11 | 2.77 | 3.18 |
| Modlow | 1.48 | 2.15 | 1.71 | 2.25 |
| Low | 0.33 | 1.18 | 0.28 | 1.14 |

**Table S2:** The range of *GC* content for the *GC* binned datasets.

|  | Autosome | X |
| --- | --- | --- |
| GC1 | [0.198, 0.556] | [0.364, 0.615] |
| GC2 | (0.556, 0.613] | (0.615, 0.68] |
| GC3 | (0.613, 0.657] | (0.68, 0.731] |
| GC4 | (0.657, 0.709] | (0.731, 0.776] |
| GC5 | (0.709, 0.963] | (0.776, 0.944] |

**Table S3:** The range of *GC* content for the *GC* binned datasets *D. simulans*.

|  | Autosome | X |
| --- | --- | --- |
| GC1 | [0, 0.564] | [0, 0.636] |
| GC2 | (0.564, 0.628] | (0.636, 0.698] |
| GC3 | (0.628, 0.674] | (0.698, 0.751] |
| GC4 | (0.674, 0.726] | (0.751, 0.799] |
| GC5 | (0.726, 0.951] | (0.799, 1] |

**Table S4:** *B* values inferred from the SFS of *GC*-changing mutations (*B*_*GC*_) (conditioned on the SFS of *GC*-conservative mutations), for autosomes, the X chromosome, and autosomes and X pooled (95% CIs constructed from likelihood ratio test are given in brackets). Estimates are either given separately for central and peripheral regions of the chromosome arms (CChr and PChr, respectively) or for whole chromosomes (WChr). All values are significantly different from *B* = 0 (*p <* 0.001). Bold values for the pooled data indicate no significant differences between autosomes and the X.

|  | WChr | CChr | PChr |
| --- | --- | --- | --- |
| Autosomes | 0.465 (0.414, 0.516) | 0.450 (0.395, 0.507) | 0.534 (0.408, 0.662) |
| X | 0.572 (0.436, 0.710) | 0.559 (0.413, 0.707) | 0.614 (0.251, 0.988) |
| Pool | **0.478 (0.430, 0.525)** | **0.464 (0.412, 0.516)** | **0.545 (0.425, 0.666)** |

**Figure S1:**
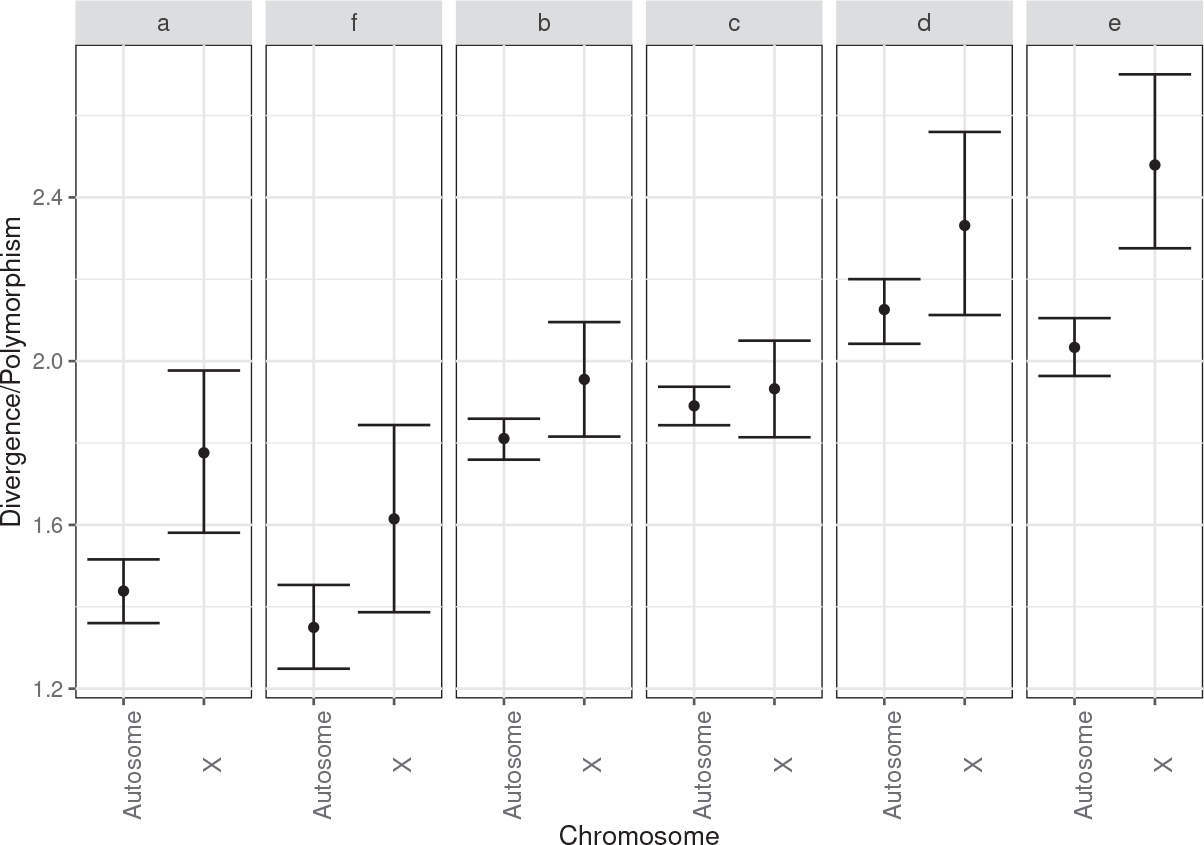
Divergence over polymorphism ratios for all six mutations classes from autosomes and X-chromosome, where *a* and *f* are *GC*-conservative, *b* and *c GC*-changing transitions and *d* and *e GC*-changing transversions.

**Figure S2:**
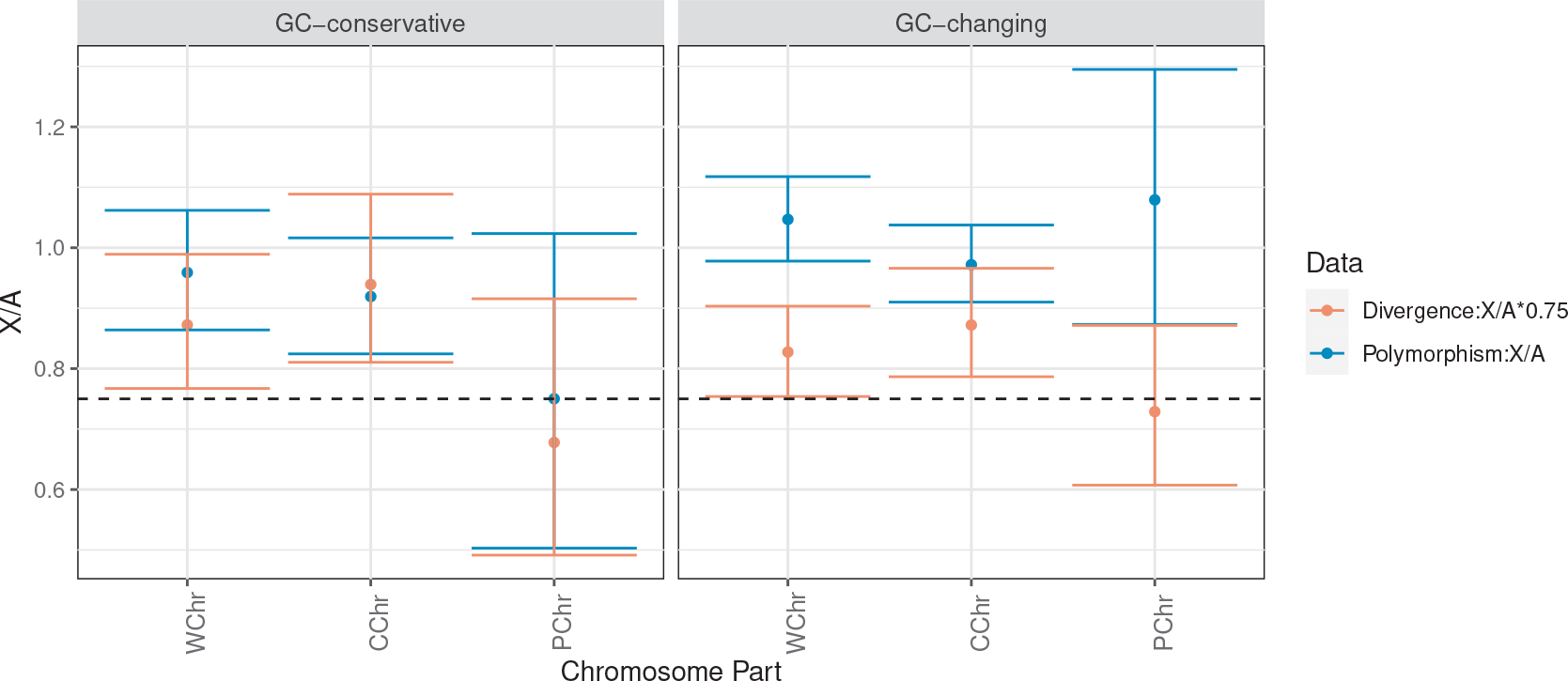
X/A ratios for polymorphism and divergence estimates given collectively for all *GC*-conservative (*a* and *f*) and changing (*b, c, d, e*) mutations. The divergence ratios are multiplied with 0.75; horizontal dashed line correspond to the value of 0.75. WChr: Whole chromosome, CChr: Central regions of the chromosome arms, PChr: Peripheral regions of the chromosome arms.

**Figure S3:**
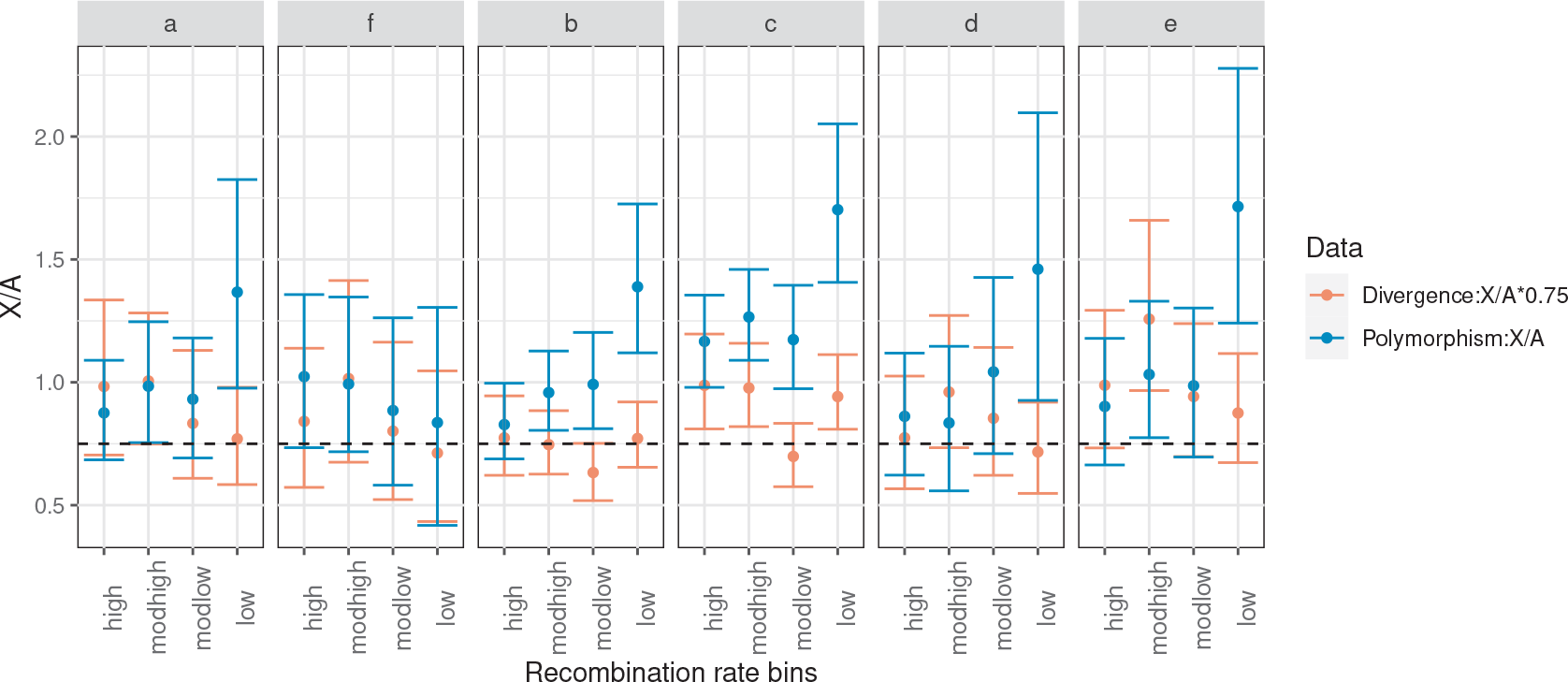
X/A ratios for polymorphism and divergence estimates for all six mutation classes; *a* and *f* are *GC*-conservative, *b* and *c GC*-changing transitions and *d* and *e GC*-changing transversions. Introns are binned by recombination rate using whole chromosome. The divergence ratios are multiplied with 0.75; horizontal dashed line correspond to the value of 0.75.

**Figure S4:**
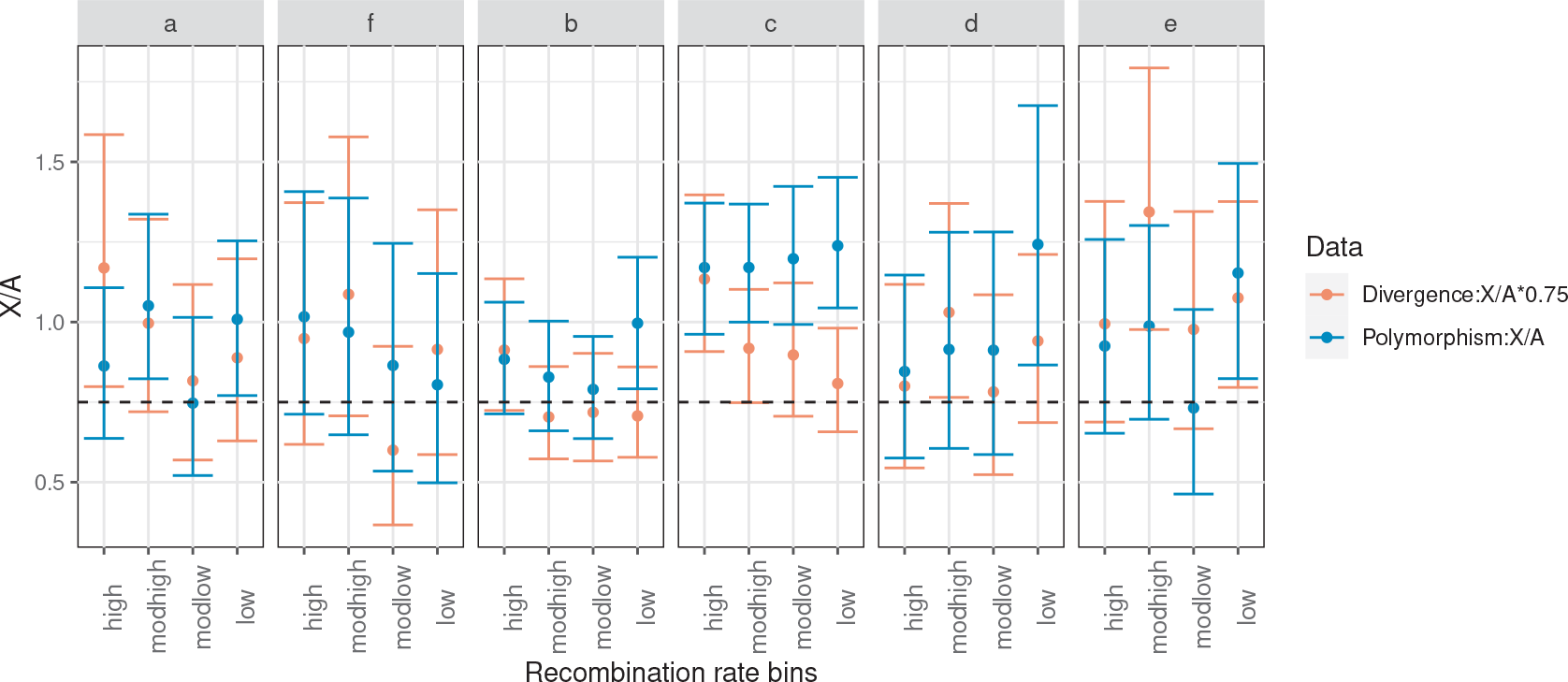
X/A ratios for polymorphism and divergence estimates for all six mutation classes; *a* and *f* are *GC*-conservative, *b* and *c GC*-changing transitions and *d* and *e GC*-changing transversions. Introns are binned by recombination rate after excluding telomeres and centromeres. The divergence ratios are multiplied with 0.75; horizontal dashed line correspond to the value of 0.75.

**Table S5:** *B* values inferred from the SFS of *GC*-changing transitions (*B*_*T s*_) and transversions (*B*_*T v*_) (conditioned on the SFS of *GC*-conservative mutations), for autosomes, the X chromosome, and autosomes and the X pooled (95% CIs constructed from likelihood ratio test are given in brackets). Estimates are either given separately for central and peripheral regions of the chromosome arms (CChr and PChr, respectively) or for whole chromosomes (WChr). All values are significantly different from *B* = 0 (*p <* 0.001). Bold values for the pooled data indicate no significant differences between autosomes and the X.

|  | WChr |  | CChr |  | PChr |  |
| --- | --- | --- | --- | --- | --- | --- |
|  | Transitions | Transversions | Transitions | Transversions | Transitions | Transversions |
| Autosome | 0.421 (0.360, 0.483) | 0.557 (0.466, 0.647) | 0.395 (0.328, 0.462) | 0.570 (0.471, 0.669) | 0.557 (0.403, 0.713) | 0.487 (0.265, 0.711) |
| X | 0.587 (0.423, 0.752) | 0.539 (0.291, 0.790) | 0.614 (0.438, 0.792) | 0.435 (0.170, 0.701) | 0.383 (-0.039, 0.816) | 1.259 (0.528, 2.08) |
| Pool | **0.441 (0.384, 0.500)** | **0.554 (0.470, 0.640)** | **0.422 (0.360, 0.486)** | **0.553 (0.461, 0.646)** | **0.540 (0.394, 0.687)** | **0.557 (0.345, 0.771)** |

**Figure S5:**
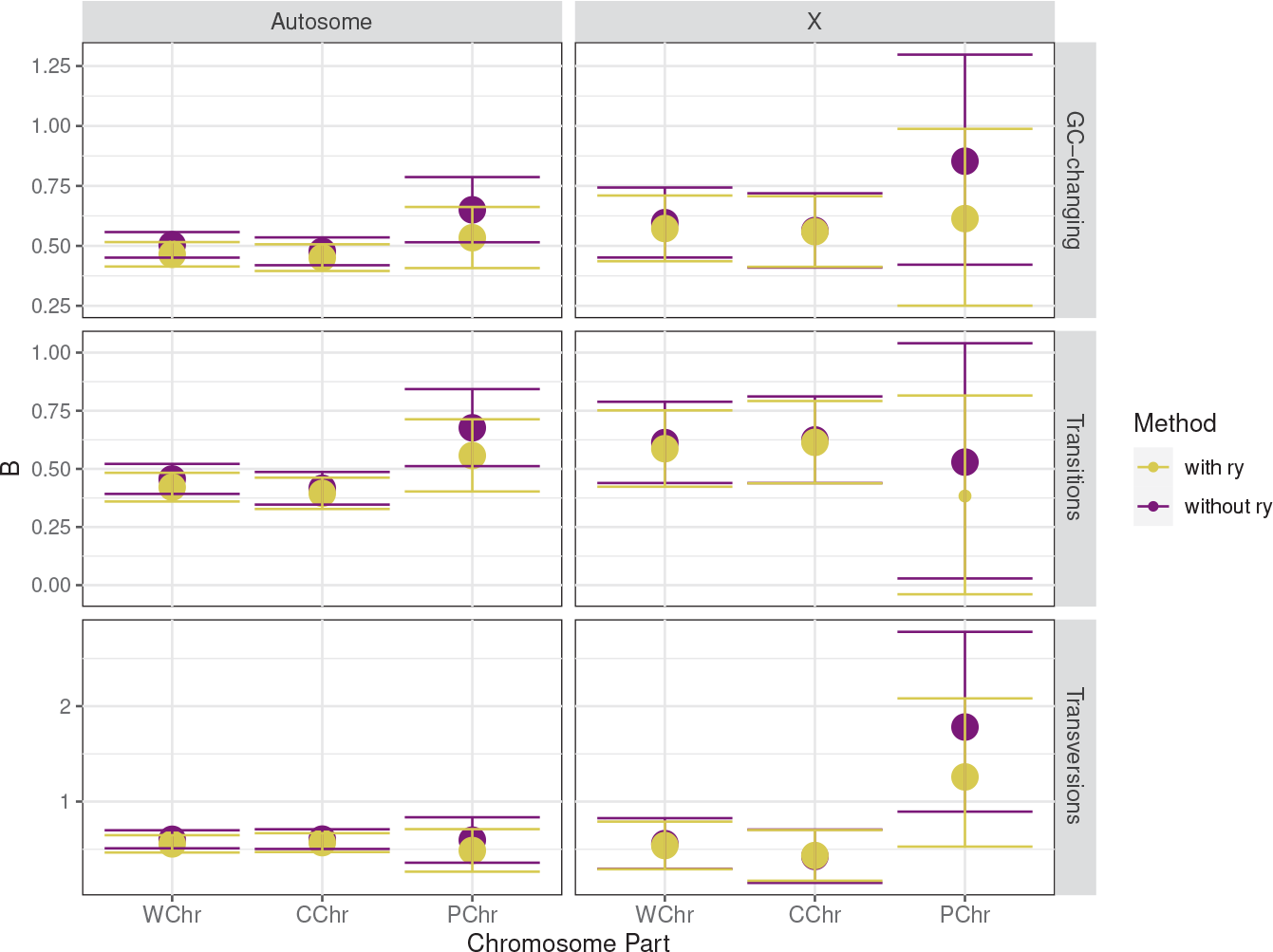
*B* values inferred from the SFS of all *GC*-changing mutations (*B*_*GC*_), *GC*-changing transitions (*B*_*Ts*_) and transversions (*B*_*Tv*_) with (yellow) or without *r*_*y*_ (purple) for autosomes and X chromosome. The big dots represents the estimates significantly different from *B* = 0. Confidence intervals are constructed from likelihood ratio test. Estimates are either given separately for central and peripheral regions of the chromosome arms (CChr and PChr, respectively) or for whole chromosomes (WChr).

**Table S6:** *B* values inferred from the SFS of all *GC*-changing mutations (*B*_*GC*_), *GC*-changing transitions (*B*_*Ts*_) and transversions (*B*_*Tv*_) (conditioned on the SFS of *GC*-conservative mutations, with *r*_*y*_) for autosomes, the X chromosome, and pooled data (autosomes+X) of *D. simulans* population (95% CIs constructed from likelihood ratio test are given in brackets). All values are significantly different from *B* = 0 (*p <* 0.001). Bold values for the pooled data indicate no significant differences between autosomes and the X.

| | $B_{GC}$ | $B_{T_v}$ | $B_{T_s}$ |
| --- | --- | --- | --- |
| Autosomes | 0.378 (0.334, 0.422) | 0.594 (0.517, 0.671) | 0.272 (0.219, 0.325) |
| X | 0.311 (0.184, 0.439) | 0.480 (0.255, 0.708) | 0.232 (0.079, 0.386) |
| Pool | **0.371 (0.330, 0.412)** | **0.582 (0.509, 0.655)** | **0.268 (0.217, 0.318)** |

**Figure S6:**
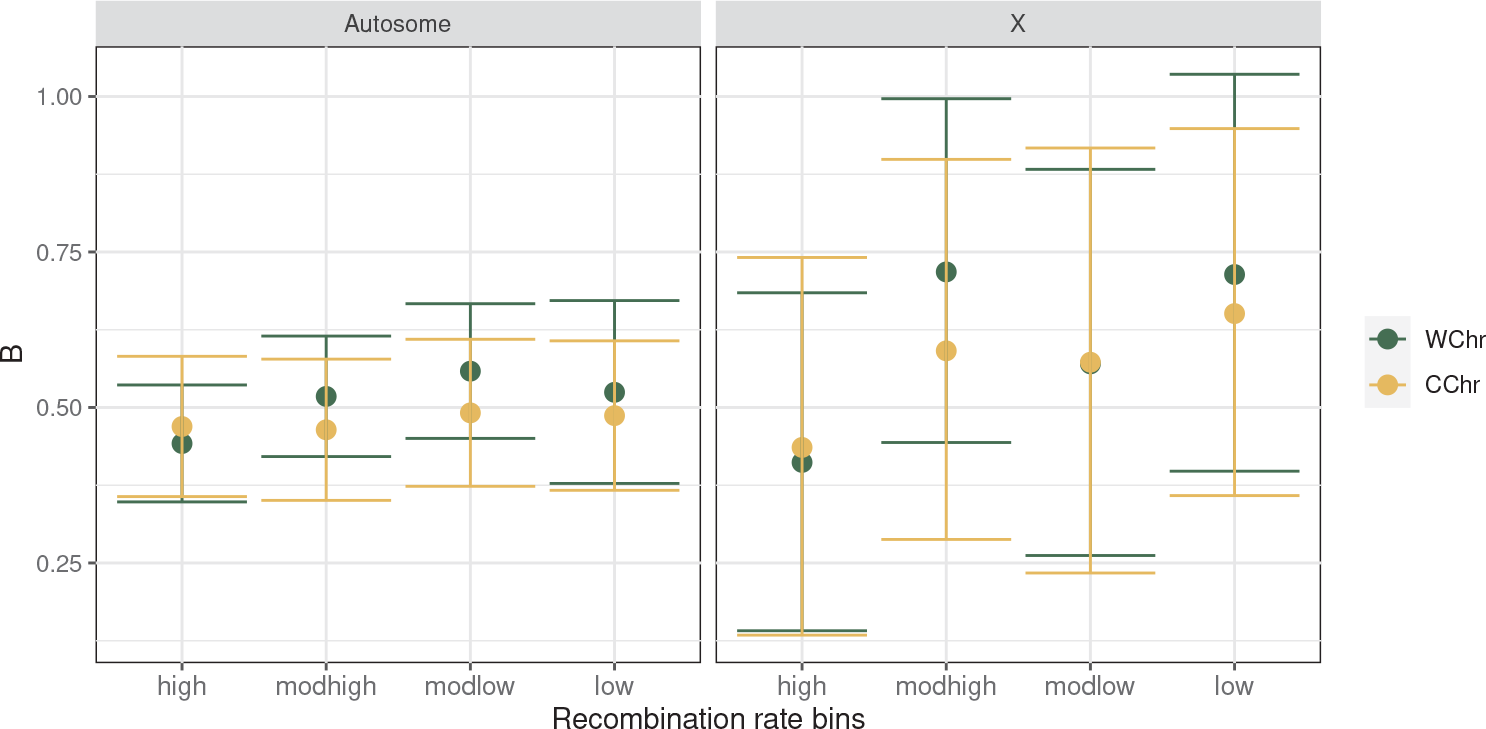
B values inferred from the SFS of *GC*-changing mutations (*B*_*GC*_) for autosomes and X chromosomes. Introns are binned by recombination rate before (green, WChr: Whole chromosome) and after excluding telomeres and centromeres (yellow, CChr: Central regions of the chromosome arms). Confidence intervals are constructed from likelihood ratio test.

**Figure S7:**
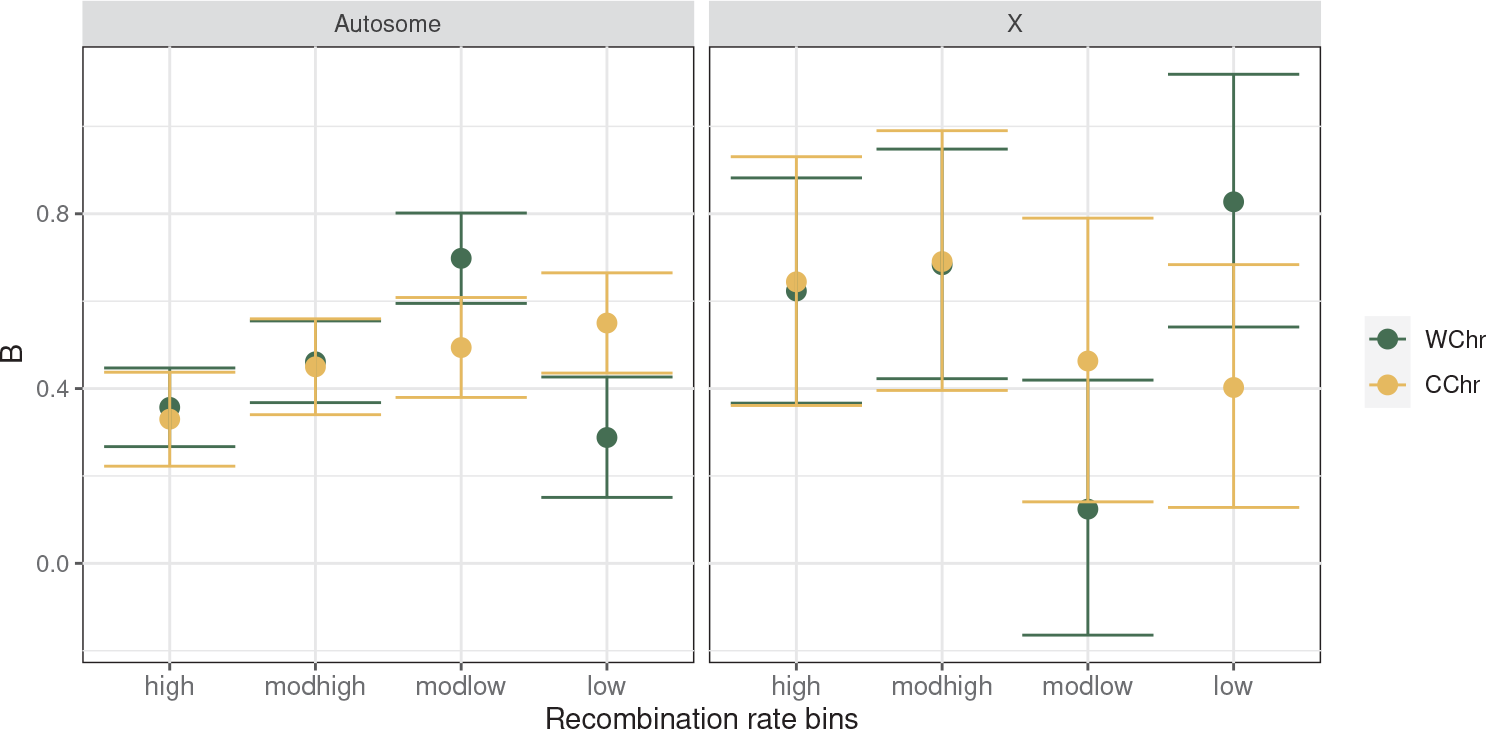
B values inferred from the SFS of *GC*-changing mutations (*B*_*GC*_) (conditioned on the SFS of *GC*-conservative mutations) for autosomes and X chromosomes. Introns are binned by recombination rate before (green, WChr: Whole chromosome) and after excluding telomeres and centromeres (yellow, CChr: Central regions of the chromosome arms). Confidence intervals are constructed from likelihood ratio test.

**Figure S8:**
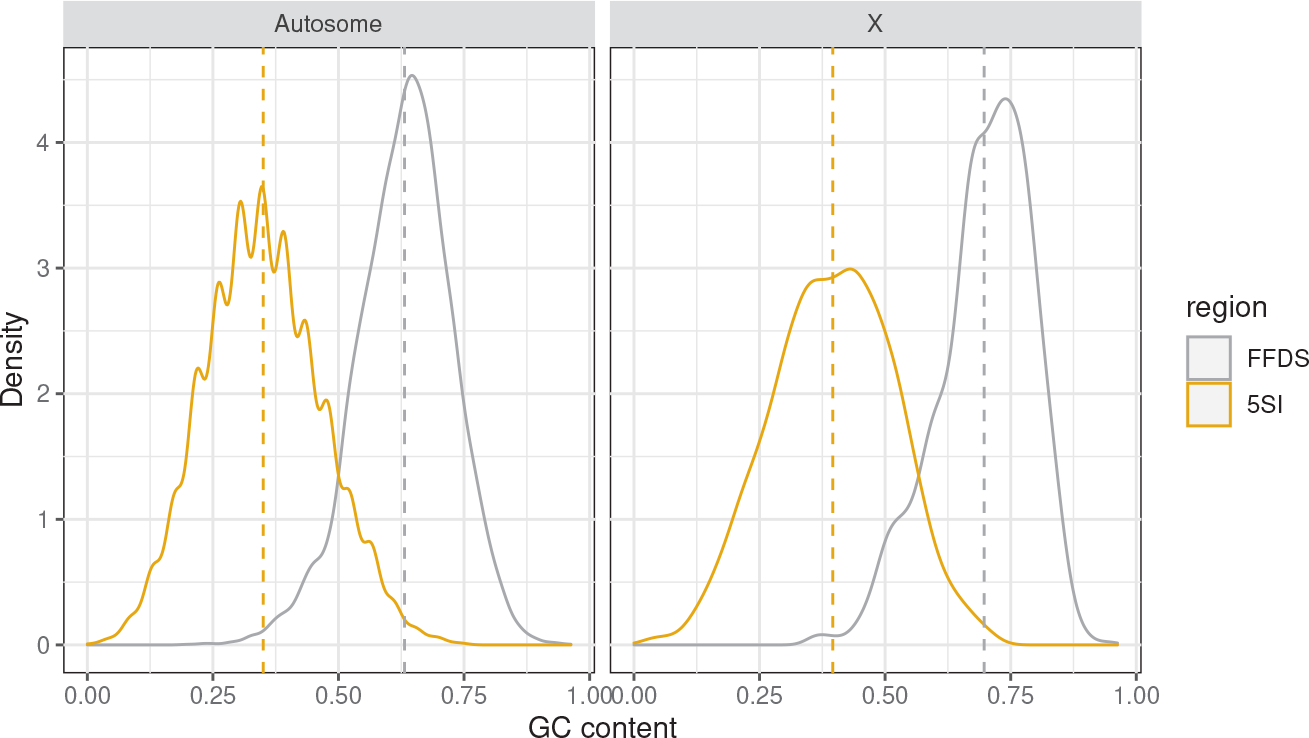
The mean *GC* content distributions for FFDS (gray) and 5SI (yellow) of autosomes and the X chromosome. The vertical dashed lines corresponds to the means of the distributions.

**Figure S9:**
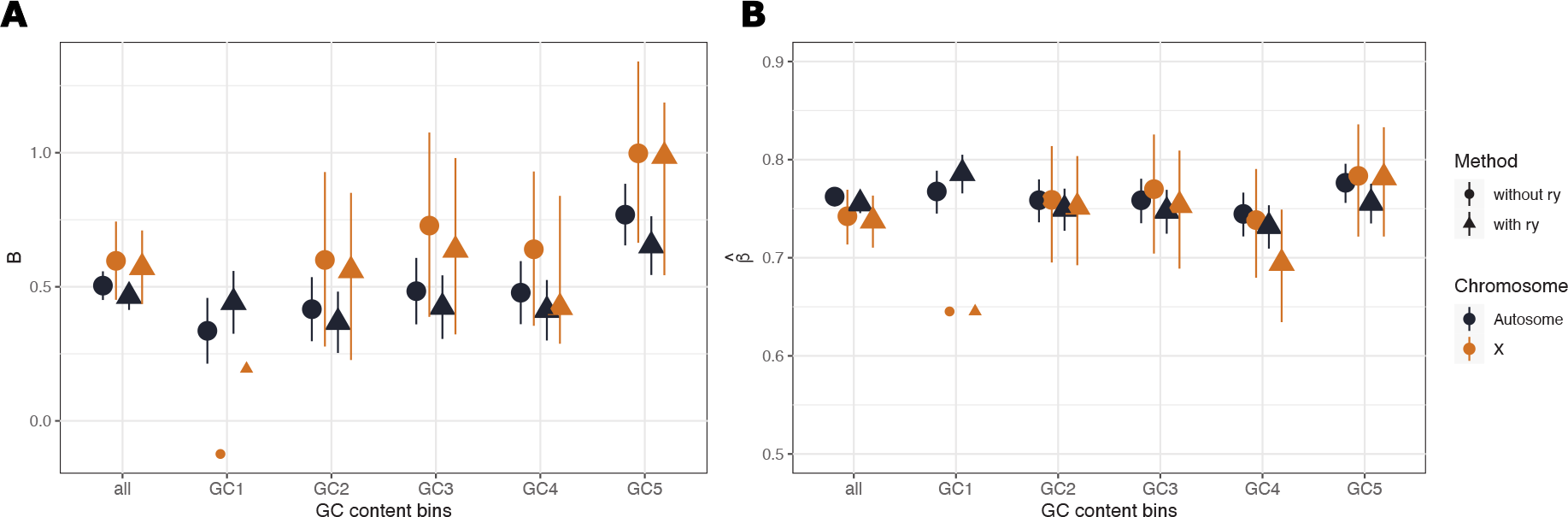
Results from *GC* content binned data of *D. melanogaster* population with (triangle) and without (round) parameter *r*_*y*_ to account for demography. A) *B*_*GC*_ values inferred from the SFS of *GC*-changing mutations for autosomes (black) and X chromosome (brown). The big dots represents the estimates significantly different from *B* = 0. Confidence intervals are constructed from likelihood ratio test. B) Mutation bias estimated conditional on *B* for for autosomes (black) and X chromosome (brown). Estimates are given for all introns and for introns binned by the mean *GC* content of the FFDS of the same genes. *GC* content increases from GC1 to GC5 and the ranges are given in S2

**Figure S10:**
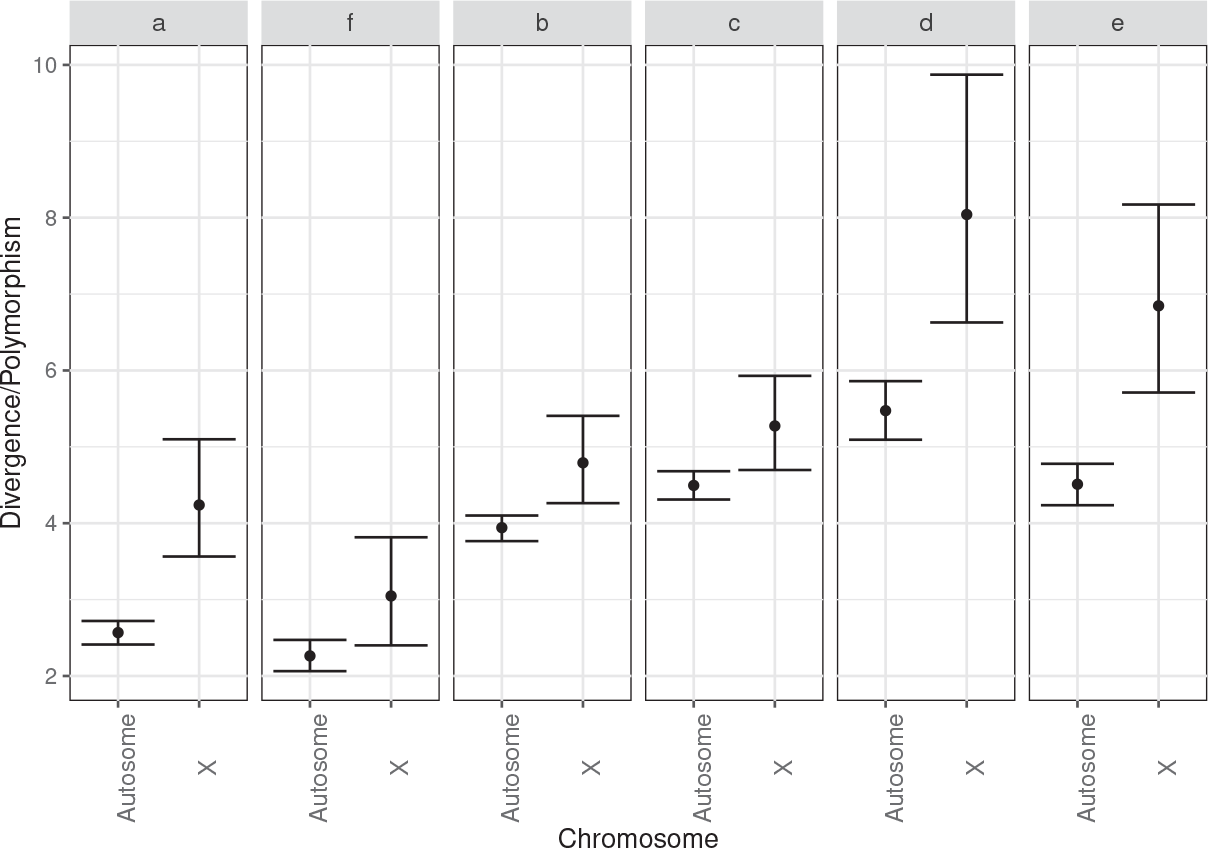
Divergence over polymorphism ratios for all six mutation classes for autosomes and the X chromosome of *D. simulans*, where *a* and *f* are *GC*-conservative, *b* and *c GC*-changing transitions and *d* and *e GC*-changing transversions.

**Figure S11:**
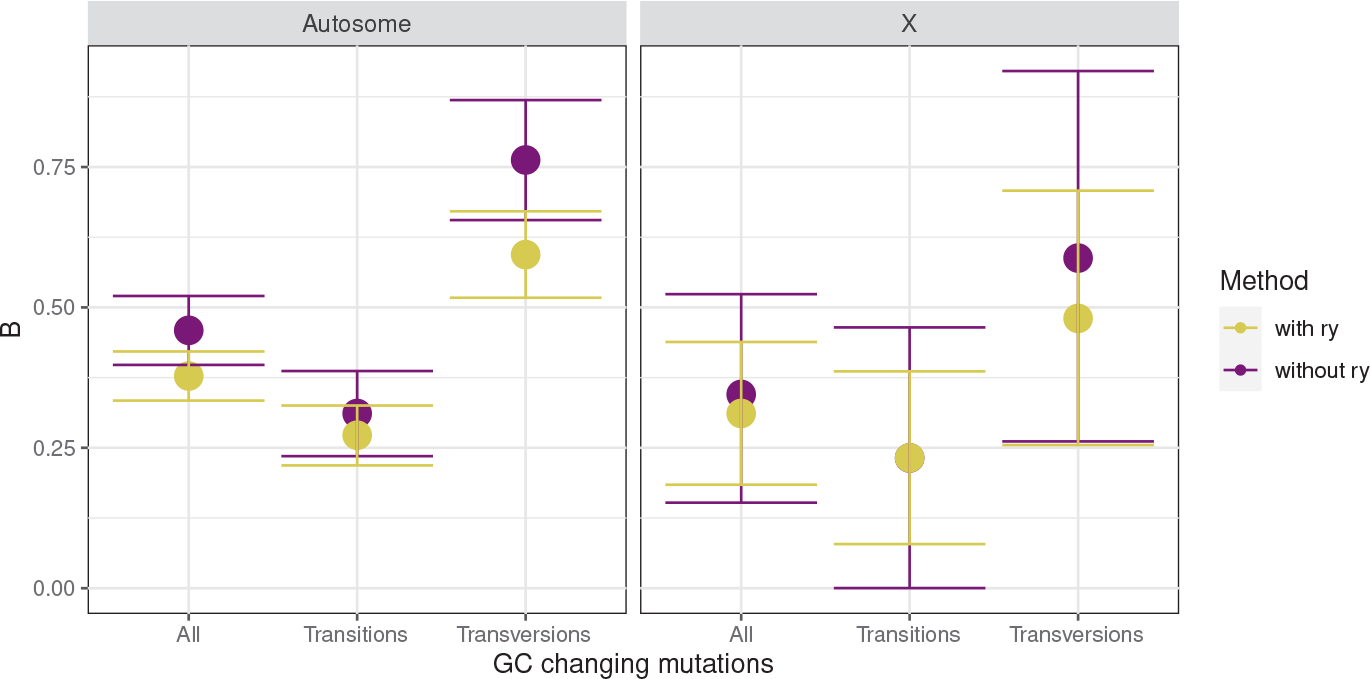
*B* values inferred from the SFS of all *GC*-changing mutations (*B*_*GC*_), *GC*-changing transitions (*B*_*Ts*_) and transversions (*B*_*Tv*_) with (yellow) or without *r*_*y*_ (purple) for autosomes and X chromosome of *D. simulans* population. The big dots represents the estimates significantly different from *B* = 0. Confidence intervals are constructed from likelihood ratio test.

**Figure S12:**
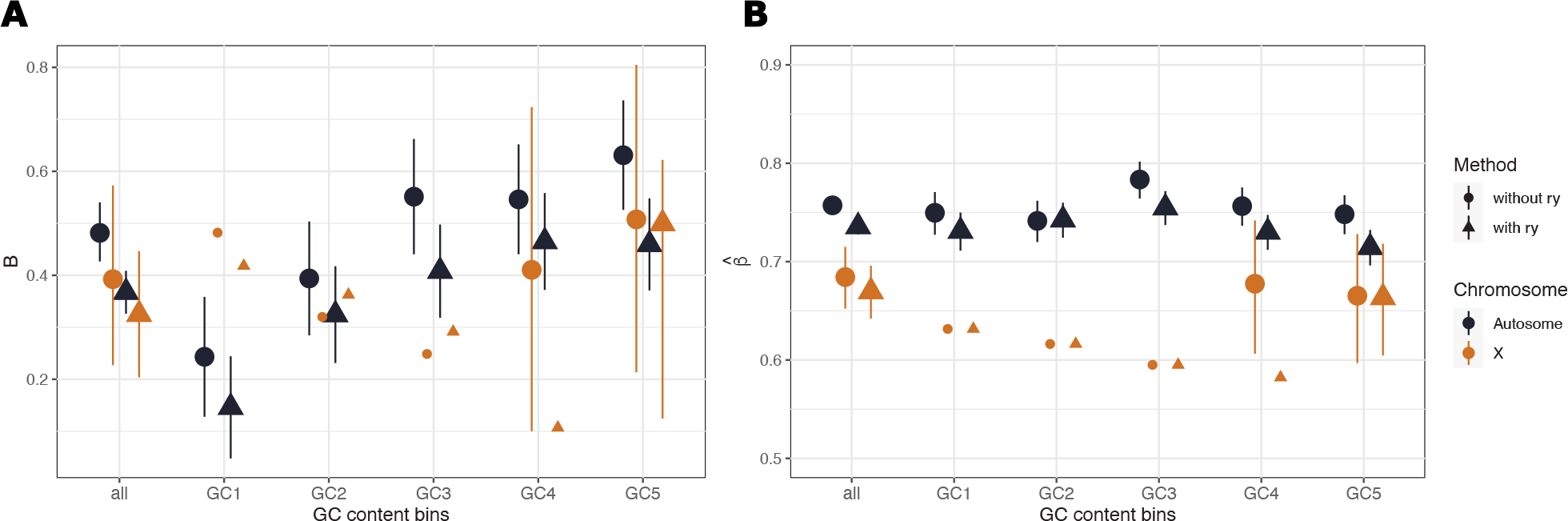
Results from *GC* content binned data of *D. simulans* population with (triangle) and without (round) parameter *r*_*y*_ to account for demography. A) *B*_*GC*_ values inferred from the SFS of *GC*-changing mutations for autosomes (black) and X chromosome (brown). The big dots represents the estimates significantly different from *B* = 0. Confidence intervals are constructed from likelihood ratio test. B) Mutation bias estimated conditional on *B* for for autosomes (black) and X chromosome (brown). Estimates are given for all introns and for introns binned by the mean *GC* content of the FFDS of the same genes. *GC* content increases from GC1 to GC5 and the ranges are given in S3

## Notes

### Competing Interest Statement

The authors have declared no competing interest.

### Summary of Updates

.

